# CRISPR screens identify targets to rescue age-related T cell dysfunction in cancer

**DOI:** 10.64898/2026.01.22.701075

**Authors:** Alex C. Y. Chen, Keely Y. Ji, Cansu Yerinde, Nelson H. Knudsen, Kevin Bi, Daniela Martinez, Thomas J. Carmona-LaSalle, Kazuhiro Taguchi, Katherine H. Xu, Elizabeth M. Seider, Marc A. Schwartz, Maria Zschummel, Linda T. Nieman, Kathleen B. Yates, Thorsten R. Mempel, Robert T. Manguso, Nir Hacohen, Debattama R. Sen

## Abstract

Immune aging is being increasingly recognized as a critical barrier to effective cancer immunotherapy, as the aged tumor microenvironment (TME) drives T cell dysfunction and impairs immune control of cancer^1–4^. However, the key molecular drivers of this process as well as potential targets to rescue T cell dysfunction in aged tumors remain incompletely understood. Therefore, we performed in vivo single-cell CRISPR screens in CD8⁺ T cells within aged tumors and tumor-draining lymph nodes (tdLNs). We identified Dusp5 and Zfp219 as key regulators of T cell persistence and effector differentiation in aged hosts. Loss of *Dusp5*, a negative regulator of ERK signaling, increased ERK1/2 phosphorylation and enhanced T cell proliferation in both young and aged tumors. In contrast, loss of *Zfp219*, a transcriptional repressor, induced epigenetic reprogramming of cytotoxic gene programs, thereby increasing granzyme secretion and enhancing antitumor immunity. Moreover, expression of the human ortholog gene *ZNF219* is increased within intratumoral CD8⁺ T cells in older cancer patients. High *ZNF219* expression correlates with poorer survival following immune checkpoint blockade (ICB) and reduces persistence of human intratumoral T cells. Notably, *Zfp219* ablation synergized with anti-PD-1 blockade in mice to expand effector-like CD8⁺ T cells, leading to significantly enhanced anti-tumor immunity and tumor clearance in aged hosts. Together, these findings highlight Dusp5 and Zfp219 as critical drivers of age-related T cell dysfunction and as potential therapeutic targets to rejuvenate T cell antitumor immunity in older cancer patients.

## Main text

As the global population continues to age, the cancer incidence among older adults is rising rapidly^3,5^. Aging is accompanied by a gradual decline in immune surveillance, which impairs the immune system’s ability to recognize and eliminate cancer cells effectively^1–4,6,7^. Among immune cells, tumor-infiltrating lymphocytes (TILs), especially CD4^+^ and CD8^+^ T cells, play vital roles in controlling cancer and are key mediators of immunotherapy effectiveness after adoptive cell therapy (ACT) or immune checkpoint inhibitor (ICI) therapy^8–11^. The antitumor activity of T cells depends heavily on their expansion, persistence, and sustained effector function. However, functional T cell immunity declines with age, leading to decreased expansion, persistence, and cytotoxic activity in the aged TME^1,2,4^. Thus, there is growing interest in understanding the factors that limit T cell control of cancer to develop strategies that can benefit more patients. While we and others have identified intrinsic defects in aged T cells, accumulating evidence indicates that extrinsic factors within the aged microenvironment further restrict T cell responses^1,12–14^. In this study, we aim to identify the genetic regulators that limit CD8^+^ T cell antitumor activity in aged tumors and to find targetable genes to reprogram T cell function and improve T cell-based cancer immunotherapy.

CRISPR-Cas9 loss-of-function screens have become powerful tools to systematically identify genetic regulators of T cell function^15–17^. Most previous screens in T cells have been conducted either in vitro or in vivo with pre-activated cells, which may not fully represent the physiological steps of naive T cell priming, differentiation, and efficacy in cancer^18^. Here, we introduce the first in vivo pooled CRISPR screen conducted in naive CD8^+^ T cells and transferred into both young or aged tumor-bearing mice. By comparing gene perturbations that improve T cell expansion, persistence, and antitumor activity between different ages, we uncover molecular pathways that govern age-related immune dysfunction and identify targets to rejuvenate T cell function in aged tumors.

### CRISPR screen identifies key regulators of T cell persistence in the aged tumor

We leveraged an in vivo single-cell CRISPR screening platform in a mouse tumor model to interrogate gene function in naïve Cas9^+^ OT-1 CD8^+^ T cells during tumor progression in aged hosts. To investigate the biology of tumor-reactive CD8^+^ T cells, we adoptively transferred OT-1 T cells into mice bearing ovalbumin (OVA)-expressing B16 melanoma (B16-OVA) tumor (Fig. 1a). We designed our lentiviral CRISPR library to target 60 transcriptional regulators and/or nuclear proteins, which were selected based on a literature review of T cell biology and differential expression analyses between young and aged T cells across distinct T cell subsets^1^. Two additional genes, *Pdcd1* and *Ptpn2*, were included as positive controls for T cell activation. To ensure statistical power, we evaluated 6 unique sgRNAs per gene, yielding a total of 412 single-guide RNAs (sgRNAs), including 20 non-targeting controls (NTCs) and 20 intergenic one-target controls (OTCs) (Extended Data Fig. 1a–e). We optimized the timing and dose of naïve Cas9^+^ OT-1 CD8^+^ T cells (2.5×10^5^ cells per mouse, one day before tumor implantation) to maximize both T cell engraftment and T cell recovery from tumors (Extended Data Fig. 1f–g). We next validated the sgRNA delivery system in vivo using Thy1.2-targeting sgRNAs and confirmed the CRISPR-Cas9 gene knockout (KO) efficiency of our platform in intratumoral Cas9^+^ OT-1 cells isolated 10 days after tumor inoculation (Extended Data Fig. 1h–i).

**Fig. 1.**
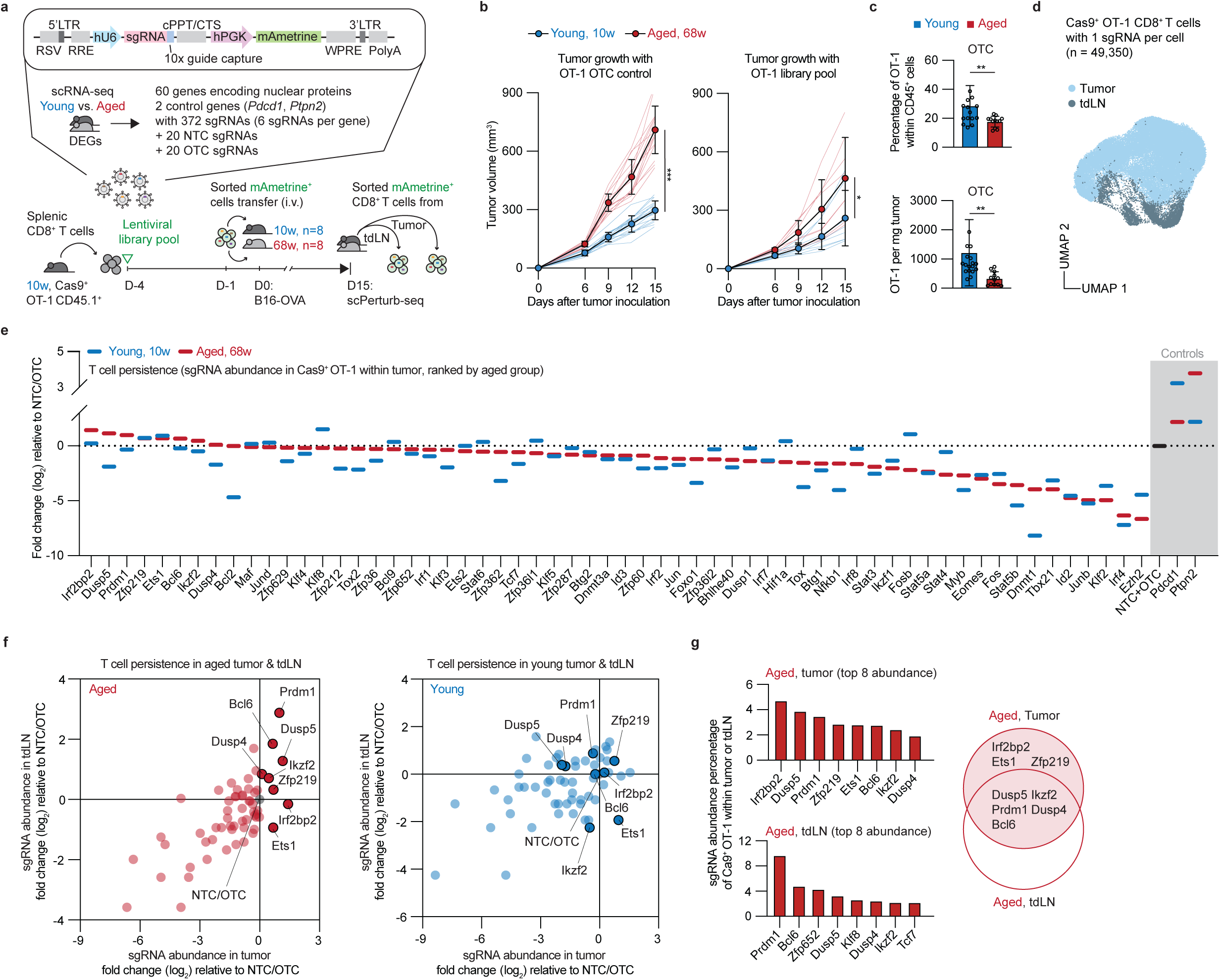
In vivo loss-of-function CRISPR screen identifies key regulators of T cell persistence in the aged tumor-bearing mice. **a**, Schema of the in vivo single-cell CRISPR screen workflow. Naive, 10-week-old Cas9^+^ OT-1 CD45.1^+^ CD8^+^ T cells were transduced with sgRNA library lentiviruses on day -4. These cells were cultured for 3 days to allow mAmetrine expression while preserving their viability and naïve state. On day -1, mAmetrine^+^ Cas9^+^ OT-1 CD8^+^ T cells were sorted, and were intravenously (i.v.) transferred into young (10-week-old, n=8) and aged (68-week-old, n=8) mice one day prior to B16-OVA tumor injection (day 0). On day 15, mice were euthanized, and mAmetrine^+^ OT-1 CD8^+^ T cells were isolated from tumors and tdLNs for Perturb-seq. **b**, Tumor growth was monitored, including mice receiving OT-1 carrying intergenic one-target controls (OTC; n = 20 young, 16 aged mice) and OT-1 carrying library pool (n = 8 young and 8 aged mice). **c**, The percentage and count of day 15 OTC OT-1 CD8^+^ T cells from B16-OVA tumors in young (10 weeks; n =18; blue) and aged (68 weeks; n =12; red) mice. **d**, Uniform manifold approximation and projection (UMAP) of Perturb-seq profiles from 49,350 OT-1 CD8^+^ T cells, each carrying a sgRNA, from tumors (aqua) and tdLNs (grey). **e**, Log2 fold change (LFC) comparison for enriched or depleted sgRNAs of each gene, relative to non-targeting controls (NTC) and OTC. Targeting genes in aged tumors is shown in red, and in young tumors in blue. **f**, LFC comparison for each targeting gene enriched in both tumors and tdLNs, relative to NTC/OTC; aged in red and young in blue. **g-h**, Top 8 enriched targeting genes in the aged tumor and tdLN, ranked by LFC. Pie chart showing the targeting genes that overlap between the aged tumor and tdLN, and those that do not. Mean ± s.d. For **b** and **c**, significance was calculated using a two-sided Student’s t-test. Asterisks used to indicate significance correspond to the following: N.S. (not significant, *P* > 0.05), \**P* ≤ 0.05, \*\**P* ≤ 0.01, and \*\*\**P* ≤ 0.001.

To perform our in vivo single-cell CRISPR screen, CRISPR library-transduced Cas9^+^ OT-1 CD45.1^+^ CD8^+^ T cells were transferred into young (10-weeks) and aged (68-weeks) CD45.2^+^ wild-type C57BL/6 mice one day prior to subcutaneous implantation of B16-OVA tumor cells. After 15 days of tumor growth, mAmetrine^+^ OT-1 T cells were sorted from tumors and tdLNs for sgRNA recovery (Extended Data Fig. 2a-d). Tumor progression was faster in aged mice, regardless of whether control or library-transduced OT-1 was transferred (Fig. 1b). Consistent with our prior findings^1^ and previous reports^2,4^, tumors in aged mice contained fewer intratumoral OT-1 T cells than young mice (Fig. 1c). To link genetic perturbations with transcriptional profiles, we performed Perturb-seq, combining CRISPR-based gene perturbation with single-cell RNA sequencing (scRNA-seq). In the end, we successfully recovered 49,350 OT-1 T cells, each carrying a sgRNA, from tumors and tdLNs (Fig. 1d).

Given that T cell expansion and persistence represent major challenges within the aged TME, we next compared sgRNA abundance between tumor and tdLN to identify genetic perturbations that enhance T cell persistence in vivo. As expected, deletion of many transcriptional regulators did not confer an advantage in T cell persistence, underscoring their essential roles in T cell maintenance, including *Dnmt1*, *Ezh2*, *Irf4*, *Junb*, and *Stat5b.* (Fig. 1e; Extended Data Fig. 2e-f; Supplementary Table 4). In contrast, sgRNAs targeting *Bcl6*, *Dusp4*, *Dusp5*, *Ets1*, *Ikzf2*, *Irf2bp2*, *Prdm1*, and *Zfp219*, along with the control genes *Pdcd1* and *Ptpn2*, were enriched in aged tumors and tdLNs, whereas only a subset of these sgRNAs was enriched in the young cohort (Fig. 1f; Supplementary Table 4). These data suggest distinct transcriptional regulators are required for T cells to adapt to aged versus young environments. Notably, *Bcl6*, *Dusp4*, *Dusp5*, *Ikzf2*, and *Prdm1* were ranked among the top enriched sgRNAs in both aged tumors and tdLNs, while *Ets1*, *Irf2bp2*, and *Zfp219* were predominantly enriched in aged tumors. These top 8 gene perturbations associated with increased T cell expansion and persistence were thus prioritized for further investigation (Fig. 1g; Supplementary Table 5).

### Loss of key T cell regulators alters subset formation in young and aged tumors

Besides the sheer number of tumor-reactive CD8^+^ T cells, emerging evidence indicates that T cell functional states, including proliferative capacity, effector function, and memory potential, are crucial determinants of T cell-mediated cancer control and are associated with improved responses to immune checkpoint blockade^9,19–22^. Thus, although the top 8 gene targets (targeting *Bcl6*, *Dusp4*, *Dusp5*, *Ets1*, *Ikzf2*, *Irf2bp2*, *Prdm1*, and *Zfp219*) promoted T cell expansion and persistence, we wondered how these perturbations might impact the differentiation state and function of T cells. To assess how these perturbations influence T cell fate decisions, we next characterized tumor-infiltrating CD8⁺ T cell subsets in tumors (Extended Data Fig. 3a-d) and tdLNs (Extended Data Fig. 3e-i) based on their transcriptional profiles. Within 43,671 CD8^+^ T cells from young and aged tumors, 38,817 T cells carried single-gene perturbations and 4,854 T cells carried with control guide RNAs, which grouped into 9 transcriptionally distinct cell clusters (Fig. 2a; Supplementary Table 6). Consistent with our previous finding^1^, the tumor-infiltrating age-associated dysfunctional (T_TAD_; high *Il7r*, *Klf2*, *Zfp36*, and *Emb*) cells were enriched in aged tumors (Fig. 2b). To link gene perturbations with single-cell transcriptional states, we examined how gene perturbations altered T cell subset formation relative to NTC/OTC sgRNAs (Extended Data Fig. 4a). Most NTC/OTC OT-1 CD8⁺ T cells were distributed among dividing (T_Divi_), progenitor-like exhausted (T_Prog_), transitory exhausted (T_Tran_), and effector-like (T_Eff-like_) cell states, with an increase in terminally exhausted (T_Term_) cell states. As anticipated, *Pdcd1* or *Ptpn2* KO directed T cells toward the T_Eff-like_ cell state. Notably, sgRNAs targeting *Bcl6*, *Ikzf2*, and *Dusp4* were predominantly localized in the T_TAD_ cluster, while *Prdm1*-KO T cells were enriched in the T_Prog_ cluster. In contrast, T cells with sgRNAs targeting *Dusp5*, *Ets1*, *Irf2bp2*, and *Zfp219* were mainly found in the T_Eff-like_ cluster (Fig. 2c). We were surprised to observe that *Dusp4* and *Dusp5* deletions drove T cells toward opposing cell states, even though *Dusp4* and *Dusp5* both belong to the same dual-specificity phosphatase (DUSP) family and are known negative regulators of the MAPK pathway.

**Fig. 2.**
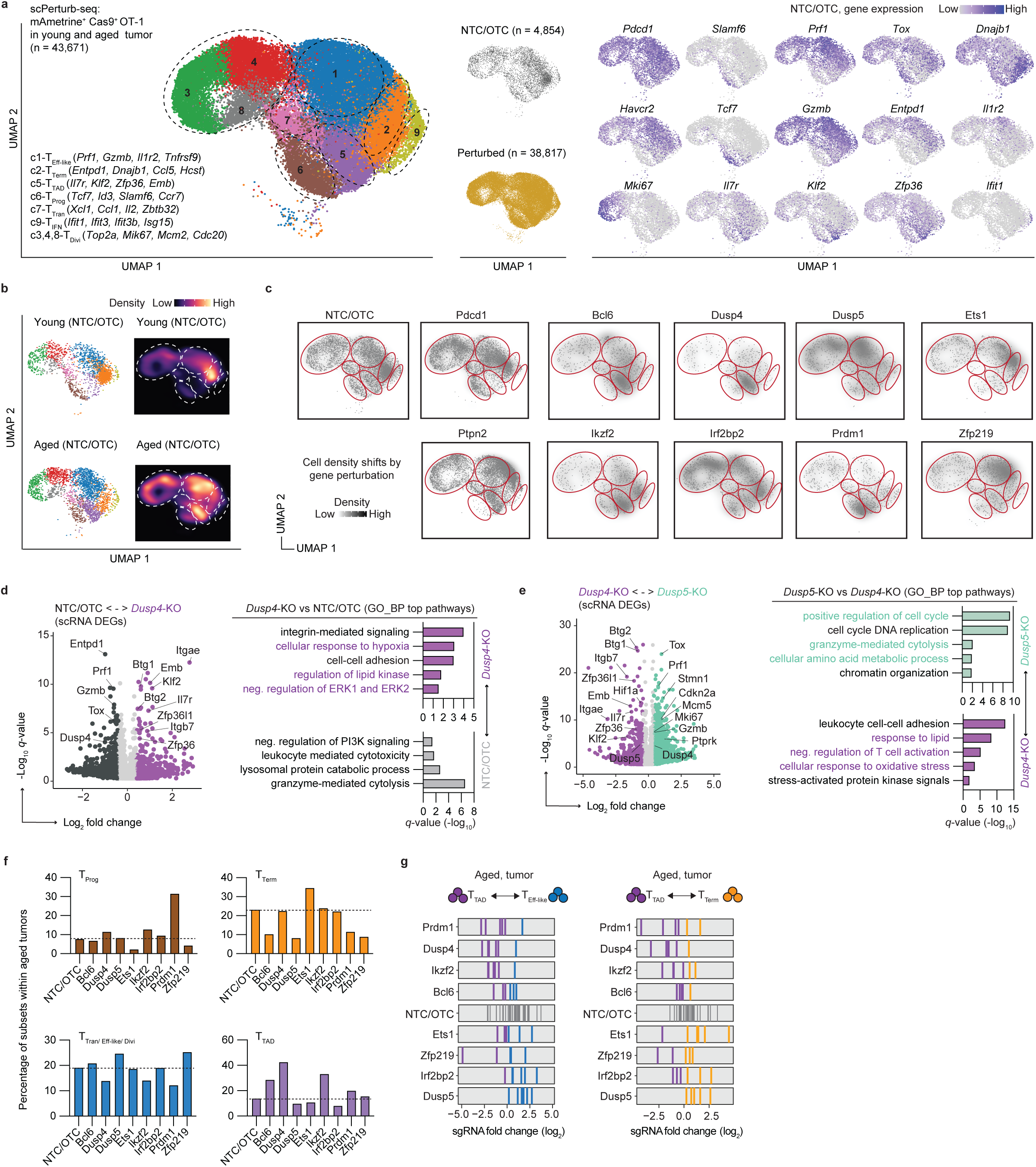
Loss of key T cell regulators alters subset formation in young and aged tumors. **a**, UMAP visualization of Perturb-seq data from 43,671 intratumoral OT-1 CD8^+^ T cells collected from day 15 B16-OVA tumors in both young and aged mice, colored by cell cluster. Of these T cells, 38,817 had single-gene perturbations, while 4,854 had NTC/OTC gRNAs. They clustered into 9 distinct transcriptional cell clusters. C1, effector-like (T_Eff-like_; blue); C2, terminally exhausted (T_Term_; orange); C5, tumor-infiltrating age-associated dysfunctional (T_TAD_; purple); C6, progenitor exhausted (T_Prog_; brown); C7, transitory exhausted (T_Tran_; pink); C9, IFN response (T_IFN_; yellow); and C3,4,8, dividing (T_Divi_) OT-1 CD8^+^ T cells. The expression levels of the specified genes are displayed on the left, with gray indicating low expression and purple indicating high expression. **b**, Galaxy plots show cell density in UMAP space for OT-1 CD8^+^ T cells from young (top) and aged (bottom) tumor-bearing mice. Cooler colors represent low density, while warmer colors indicate high density. **c**, Cell density projections for T cell clusters and subset shifts in tumors following gene perturbation. Lighter shades represent low cell density, whereas darker shades indicate high cell density. **d-e**, Volcano plot showing differentially expressed genes (DEGs) among transferred OT-1 CD8^+^ T cells in tumors: NTC/OTC (dark grey), *Dusp4* knockout (*Dusp4*-KO, purple), and *Dusp5*-KO (green). GO biological process pathway analysis of DEGs between groups is included, with each q-value (-log10). **f**, Percentage of subsets within aged tumors for each gene perturbation. **g**, Frequency histograms show enrichment of sgRNAs for T_TAD_ (purple), T_Eff-like_ (blue), and T_Term_ (orange). The dashed black line indicates the average LFC of NTC/OTC.

We hypothesized that *Dusp4* and *Dusp5* play distinct regulatory roles within the broader DUSP family, and so we next assessed whether perturbing these genes would have a differential impact on key gene programs including mitogen-activated protein kinase (MAPK) signaling. Compared to NTC/OTC or *Dusp5*-KO, *Dusp4*-KO T cells exhibited a significant enrichment of cellular stress responses and negative regulation of T cell activation, including high expression of T cell quiescence genes (*Btg1* and *Btg2*)^23^ and age-associated dysfunctional genes (*Il7r*, *Klf2*, *Zfp36*, and *Zfp36l1*)^1^. By contrast, *Dusp5*-KO T cells displayed elevated cell cycle (*Stmn1*, *Cdkn2a*, *Mki67*, and *Mcm5*) and granzyme-mediated cytolysis (*Gzmb* and *Prf1*) genes, consistent with functional pathway enrichment analyses (Fig. 2d-e; Supplementary Tables 7 and 8). Immune response analysis^24^ revealed that *Dusp4*-KO T cells had increased IL-17 and decreased IL-2/IL-15 response signatures, whereas *Dusp5*-KO T cells showed the opposite pattern, aligning with their proliferation and effector gene signatures (Extended Data Fig. 5a). MAPK gene enrichment further indicated that *Dusp4*-KO T cells shifted toward JNK and ERK5 pathways, whereas *Dusp5*-KO cells predominantly engaged ERK1/2 pathway (Extended Data Fig. 5b-c).

Taken together, these analyses suggest that, among the 8 prioritized gene perturbations, targeting *Dusp5*, *Ets1*, *Irf2bp2*, and *Zfp219* preferentially drive CD8⁺ T cells toward effector-like states, whereas *Dusp4*, *Prdm1*, *Ikzf2*, and *Bcl6* do not (Fig. 2f-g, Supplementary Table 9).

### *Dusp5* and *Zfp219* KO leads to increased tumor control and effector-like T cell formation

Having identified perturbations that favor effector-like T cell differentiation, we next asked whether these gene (*Dusp5*, *Ets1*, *Irf2bp2*, and *Zfp219*) deletions could rescue age-associated T cell dysfunction and enhance antitumor responses in vivo. To assess this, we transferred Cas9^+^ OT-1 CD45.1^+^ CD8^+^ T cells carrying each gene deletion into young and aged B16-OVA tumor-bearing mice (Fig. 3a). Mice that received *Dusp*5-KO and *Zfp219*-KO T cells exhibited improved tumor control in aged mice, whereas *Ets1*-KO and *Irf2bp2*-KO T cells showed similar tumor growth to OTC. Notably, loss of *Dusp5* enhanced tumor control in both young and aged mice, whereas the antitumor benefit of *Zfp219* loss was restricted to aged hosts (Fig. 3b-c).

**Fig. 3.**
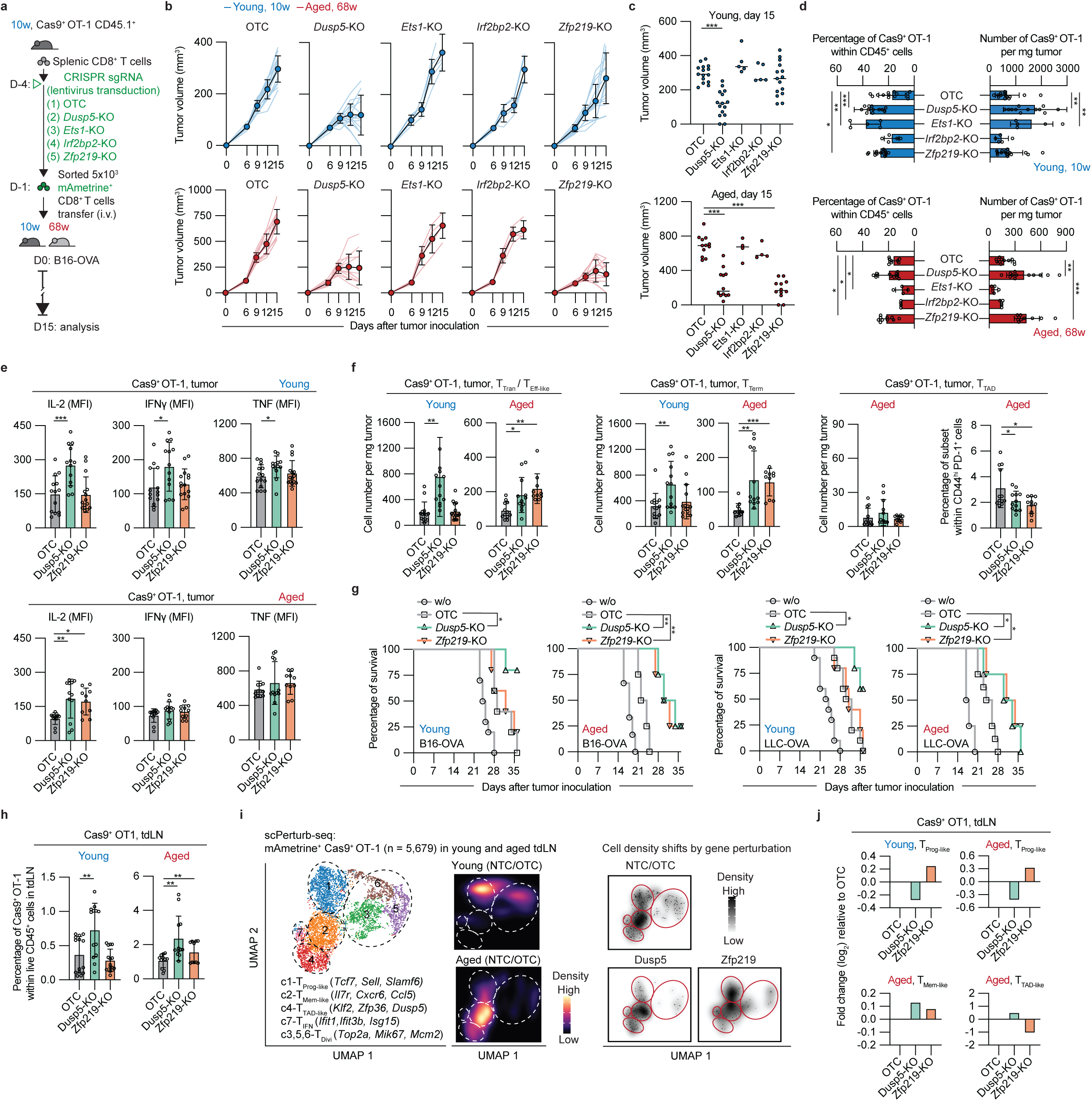
*Dusp5* and *Zfp219* KO leads to increased tumor control and effector-like T cell formation. **a-c**, Schema of the experimental design for the B16-OVA melanoma model used to evaluate tumor growth in mice receiving each gene knockout (KO) OT-1 CD8^+^ T cells. Tumor growth curves for B16-OVA tumors in young (10 weeks; n = 5-15; blue) and aged (68 weeks; n = 4-12; red) mice. The final B16-OVA tumor volumes were measured on day 15. **d**, The percentage and count of day 15 OT-1 CD8^+^ T cells after gene perturbation in tumors of young (10 weeks; n = 5-15; blue) and aged (68 weeks; n = 4-12; red) B16-OVA tumor-bearing mice. **e**, Geometric mean fluorescence intensity (MFI) of IL-2, IFNγ, and TNF in intratumoral OT-1 CD8^+^ T cells after gene perturbation (e.g., Dusp5-KO and Zfp219-KO) in young (n = 13-15; blue) and aged (n = 10-12; red) tumor-bearing mice. **f**, The count and percentage of day 15 OT-1 CD8^+^ T cell subsets after gene perturbation in young (n = 12-15; blue) and aged (n = 10-12; red) mice. **g**, Overall survival of B16-OVA- or LLC-OVA young (n = 5-10) and aged (n = 4-6) tumor-bearing mice treated with OTC, *Dusp5*-KO, or *Zpf219*-KO OT-1 CD8^+^ T cells. **h**, The percentage of day 15 OT-1 CD8^+^ T cells after gene perturbation in tdLNs of young (10 weeks; n = 13-15; blue) and aged (68 weeks; n = 10-12; red) B16-OVA tumor-bearing mice. **i**, UMAP visualization of Perturb-seq data from 5,679 OT-1 CD8^+^ T cells collected from day 15 B16-OVA tdLNs in both young and aged mice, colored by cell cluster. They clustered into 7 distinct transcriptional cell clusters. C1, progenitor-like exhausted (T_Prog-like_; blue); C2, memory-like (T_Mem-like_; orange); C4, tumor-infiltrating age-associated dysfunctional-like (T_TAD-like_; red); C7, IFN response (T_IFN_; pink); C3,5,6, dividing (T_Divi_) OT-1 CD8^+^ T cells. Galaxy plots show cell density in UMAP space for OTC OT-1 CD8^+^ T cells from young (top) and aged (bottom) tumor-bearing mice. Cooler colors represent low density, while warmer colors indicate high density. Cell density projections (right side) for T cell clusters and subset shifts in tdLNs following gene perturbation. Lighter shades represent low cell density, whereas darker shades indicate high cell density. **j**, LFC comparison for each targeting gene enriched in different T cell subsets in tdLN, relative to NTC/OTC. Mean ± s.d. For **c-f** and h, significance was calculated using a two-sided Student’s t-test. For **g**, significance was measured by the log-rank (Mantel–Cox) test. Asterisks used to indicate significance correspond to the following: N.S. (not significant, *P* > 0.05), \**P* ≤ 0.05, \*\**P* ≤ 0.01, and \*\*\**P* ≤ 0.001.

In line with their tumor control results, we observed an increased number of *Dusp5*-KO and *Zfp219*-KO T cells in aged tumors, compared to OTC (Fig. 3d and Extended Data Fig. 6a-b). These expanded T cells in aged tumors exhibited higher levels of IL-2, while IFN-γ and TNF levels remained similar to OTC (Fig. 3e). To validate our Perturb-seq results, we performed flow cytometry and confirmed that *Dusp5*-KO and *Zfp219*-KO T cells shifted toward a more T_Eff-like_ or T_Term_ cell states and showed reduced T_TAD_ formation in aged tumors (Fig. 3f and Extended Data Fig. 6c). To validate tumor control in another tumor model, we included the Lewis lung carcinoma (LLC)-OVA model. Mice bearing B16-OVA and LLC-OVA tumors showed prolonged tumor control and improved survival in aged mice receiving *Dusp5*-KO and *Zfp219*-KO T cells compared with controls (Fig. 3g).

The prolonged tumor control and survival seen in mice treated with *Dusp5*-KO or *Zfp219*-KO T cells led us to ask whether these gene perturbations also promote the persistence of tumor-specific T cells in tdLNs, which are known to act as reservoirs that support long-term antitumor immunity^25,26^. In aged mice, both *Dusp5*-KO and *Zfp219*-KO T cells were present at higher frequencies in tdLNs (Fig. 3h). Perturb-seq analysis of tdLN T cells revealed that *Dusp5*-KO T cells preferentially adopted a memory-like (T_Mem-like_) cell state, while *Zfp219*-KO T cells shifted toward both progenitor-like (T_Prog-like_) and T_Mem-like_ cell states (Fig. 3i-j; Supplementary Table 10).

### *Dusp5* KO boosts T cell proliferation while *Zfp219* KO reprograms T cell cytotoxic programs

Our findings indicate that *Dusp5* and *Zfp219* are key regulators of age-related T cell dysfunction and could serve as potential targets for restoring T cell function. However, the precise mechanisms underlying their effects remain unclear. We first examined whether the expression levels of *Dusp5* (human ortholog *DUSP5*) and *Zfp219* (human ortholog *ZNF219*) in human T cells vary with age. We reanalyzed human tumor-infiltrating CD8^+^ T cells using published pan-cancer scRNA-seq datasets^8,9,27,28^, and found comparable *DUSP5* levels across age groups, whereas *ZNF219* levels increased with age in cancer patients (Fig. 4a). Consistently, in our mouse cohort, *Zfp219* levels were elevated in T cells from aged hosts, particularly when young T cells were transferred into aged tumor-bearing mice. In contrast, aged T cells transferred into young hosts did not show this increase, suggesting that the aged environment likely drives *Zfp219* upregulation, which may explain why improved antitumor responses following Zfp219 deletion were observed only in aged mice (Fig. 4b).

**Fig. 4.**
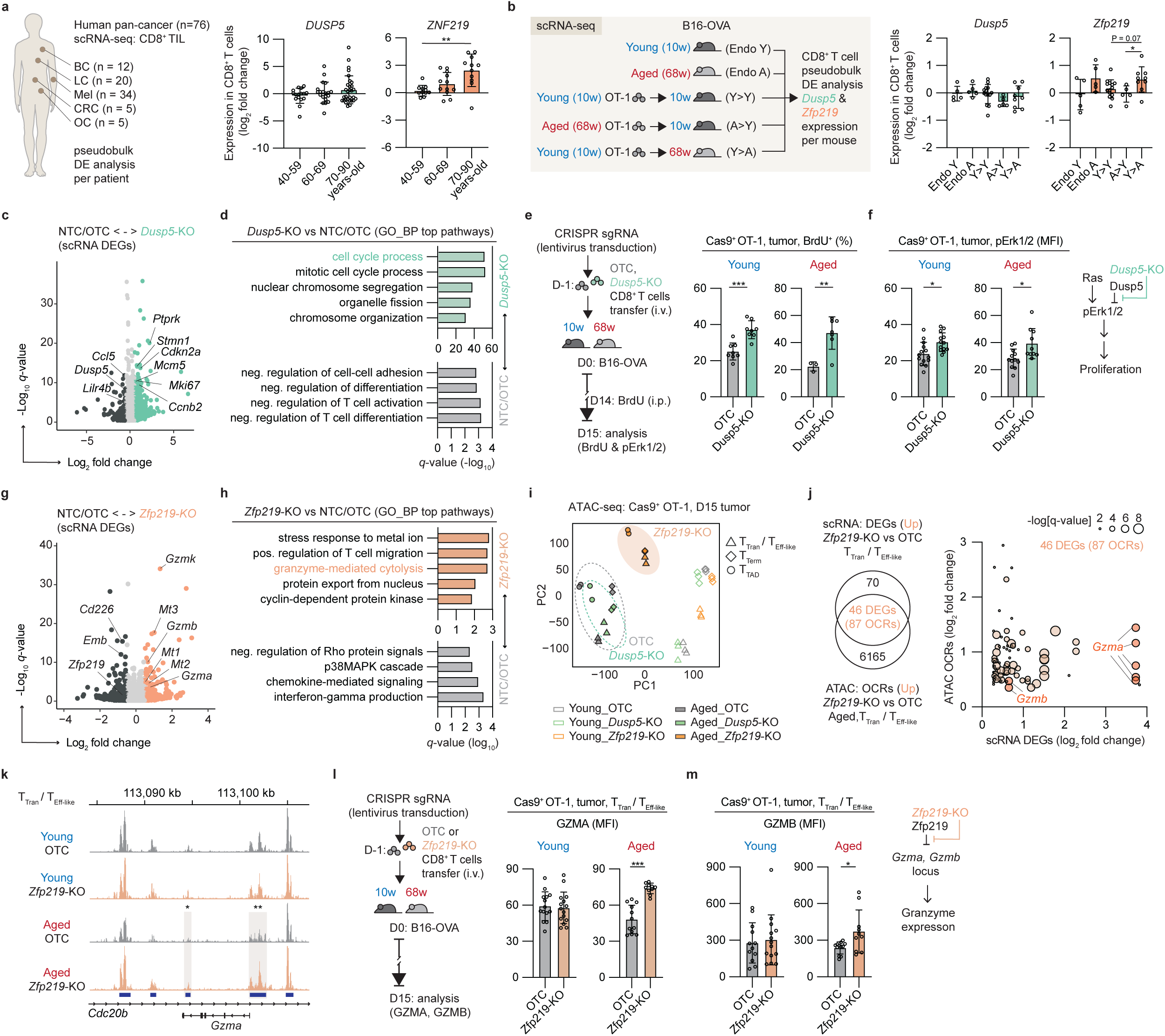
*Dusp5* KO boosts T cell proliferation while *Zfp219* KO reprograms T cell cytotoxic programs in aged tumors. **a**, Human pan-cancer scRNA-seq dataset of intratumoral CD8^+^ T cells; BC, breast cancer; LC, lung cancer; Mel, melanoma; CRC, colorectal cancer; OC, ovarian cancer. Raw count data from the scRNA-seq dataset were used to perform pseudobulk differential expression analysis. LFC comparison of *DUSP5* and *ZNF219* expression across age groups (40-50, 60-69, and 70-90 years old). **b**, scRNA-seq dataset of intratumoral endogenous CD8^+^ T cells from young (endo Y) or aged (endo aged) mice, the intratumoral transferred young OT-1CD8^+^ T cells to young (Y>Y) or aged (Y>A) mice, and the intratumoral transferred aged OT-1CD8^+^ T cells to young (A>Y) mice. LFC comparison of *Dusp5* and *Zfp219* expression across different groups. **c-d**, Volcano plot showing DEGs among transferred OT-1 CD8^+^ T cells in tumors: NTC/OTC (dark grey) and *Dusp5*-KO (green). GO biological process pathway analysis of DEGs between groups is included, with each *q*-value (-log10). **e-f**, Schema for in vivo validation of *Dusp5*-KO OT-1 CD8^+^ T cells’ proliferative capacity via flow cytometry assays measuring BrdU incorporation and Erk1/2 phosphorylation. It includes the percentage of BrdU-labeled (BrdU^+^) T cells and the MFI of phosphorylated Erk1/2 (pErk1/2) in *Dusp5*-KO T cells from young (10 weeks; n = 8-14) and aged (68 weeks; n = 3-12) B16-OCA tumor bearing mice. **g-h**, Volcano plot showing DEGs among transferred OT-1 CD8^+^ T cells in tumors: NTC/OTC (dark grey) and *Zfp219*-KO (orange). GO biological process pathway analysis of DEGs between groups is included, with each *q*-value (-log10). **i**, Principal component analysis (PCA) projection of chromatin accessibility profiles from tumor-infiltrating OT-1 T cell subsets. **j**, Overlapping enrichment observed across 87 open chromatin regions (OCRs) and 46 differentially expressed genes (DEGs) in *Zfp219*-KO compared to OTC. The scatter plot illustrates the LFC comparison, highlighting enrichment in chromatin accessibility and gene expression relative to OTC; circle size indicates the -log(*q*-value). **k**, Representative ATAC-seq tracks showing OTC (gray) and *Zfp219*-KO (orange) OT-1 cells at *Gzma* loci. **l-m**, Schema for in vivo validation of *Zfp219*-KO OT-1 CD8^+^ T cells’ granzyme secretion using flow cytometry. It includes the MFI of GZMA and GZMB in *Zfp219*-KO T cells from young (10 weeks; n = 12-15) and aged (68 weeks; n = 10-12) B16-OVA tumor-bearing mice. Mean ± s.d. For **a-b**, **e-f**, **k**, and **l-m**, significance was calculated using a two-sided Student’s t-test. Asterisks used to indicate significance correspond to the following: N.S. (not significant, *P* > 0.05), \**P* ≤ 0.05, \*\**P* ≤ 0.01, and \*\*\**P* ≤ 0.001.

Gene expression and pathway analyses further revealed that cell-cycle signatures were selectively upregulated in *Dusp5*-KO T cells (Fig. 4c-d; Supplementary Table 11). As reported previously^29,30^, Dusp5 functions as a negative feedback regulator that dephosphorylates and inactivates Erk1/2 signals, thereby limiting T cell proliferation. To validate this mechanism in vivo, we transferred *Dusp5*-KO Cas9^+^ OT-1 CD45.1^+^ CD8^+^ T cells into young and aged B16-OVA tumor-bearing mice and evaluated BrdU incorporation and Erk1/2 phosphorylation. After 15 days of tumor growth, the proportion of BrdU^+^ dividing T cells was increased in *Dusp5*-KO T cells (Fig. 4e), which also exhibited higher levels of phosphorylated Erk1/2 (pErk1/2, Fig. 4f), confirming enhanced proliferative capacity.

Conversely, *Zfp219*-KO T cells showed enriched signatures of granzyme-mediated cytolysis (Fig. 4g-h; Supplementary Table 12). As a DNA-binding transcriptional repressor^31,32^, Zfp219 may regulate cytotoxic gene expression at the epigenetic level. To verify this, we profiled chromatin accessibility in T cell subsets from young and aged B16-OVA tumor-bearing mice using ATAC-seq. *Zfp219*-KO T cells from aged tumors exhibited a distinct epigenetic landscape compared with young *Zfp219*-KO T cells, as well as with *Dusp5*-KO and OTC T cell controls. Integrative analysis of open chromatin regions (OCRs) and differentially expressed genes (DEGs) revealed overlapping enrichment of *Gzma* and *Gzmb* across 87 OCRs and 46 DEGs (Fig. 4i-j and Extended Data Fig. 7a-c). Altered chromatin accessibility at the *Gzma* locus was observed in aged, but not young, *Zfp219*-KO T cells relative to OTC controls (Fig. 4k and Extended Data Fig. 7d). Functional validation via adoptive transfer of *Zfp219*-KO T cells confirmed that, while young OTC and *Zfp219*-KO T cells expressed similar levels of GZMA and GZMB, aged OTC T cells showed a significant reduction. Importantly, deletion of *Zfp219* in T cells restored GZMA and GZMB expression in aged tumors (Fig. 4l-m and Extended Data Fig. 7e-f).

Together, these findings indicate that *Dusp5*-KO and *Zfp219*-KO enhance antitumor immunity through distinct mechanisms. Loss of *Dusp5*, a negative regulator of ERK signaling, enhances ERK1/2 phosphorylation and promotes T cell proliferation in both young and aged tumors. In contrast, loss of *Zfp219* drives epigenetic reprogramming of cytotoxic gene loci, increasing granzyme production and improving antitumor responses specifically in aged T cells (Extended Data Fig. 7g).

### *Zfp219* KO enhances responses to immune checkpoint blockade in aged hosts

The above findings, especially the age-related increase in *Zfp219* (human *ZNF219*) levels observed in intratumoral CD8⁺ T cells of older cancer patients (Fig. 4a), raise the possibility that targeting Zfp219 in T cells could be a therapeutic strategy to rejuvenate aged antitumor immunity and improve responses to immune checkpoint blockade. First, to assess the clinical relevance of our findings, we reanalyzed publicly available human pan-cancer datasets^33^, including cohorts treated with anti-PD-1 therapy. Elevated ZNF219 expression in CD8^+^ T cells was significantly associated with poorer overall survival (OS) and progression-free survival (PFS) in pretreatment (pre-tx) tumor biopsy samples (OS hazard ratio = 2.02, 95% CI = 1.49–2.75, P = 4.8 × 10^-6^; PFS hazard ratio = 2.58, 95% CI = 1.87–3.54, P = 1.5 × 10^-9^). Similar associations were observed in biopsies obtained during or after anti-PD-1 treatment (anti-PD-1 tx, OS hazard ratio = 2.53, 95% CI = 1.36–4.73, P = 0.0026; PFS hazard ratio = 3.52, 95% CI = 0.95–13.05, P = 0.045; Fig. 5a). To verify the functional impact of elevated ZNF219 expression in human T cells, we overexpressed *ZNF219* in primary human T cells from health donors and performed an in vivo T cell persistence assay. In this assay, human T cells were transferred into immunodeficient mice with established A375 melanoma tumors engineered to express anti-CD3 scFV and CD80 as an TCR-independent tumor targeting mechanism. We observed that *ZNF219* overexpression impaired T cell persistence and expansion in vivo but not in vitro relative to the mCherry overexpression control (Fig. 5b).

**Fig. 5.**
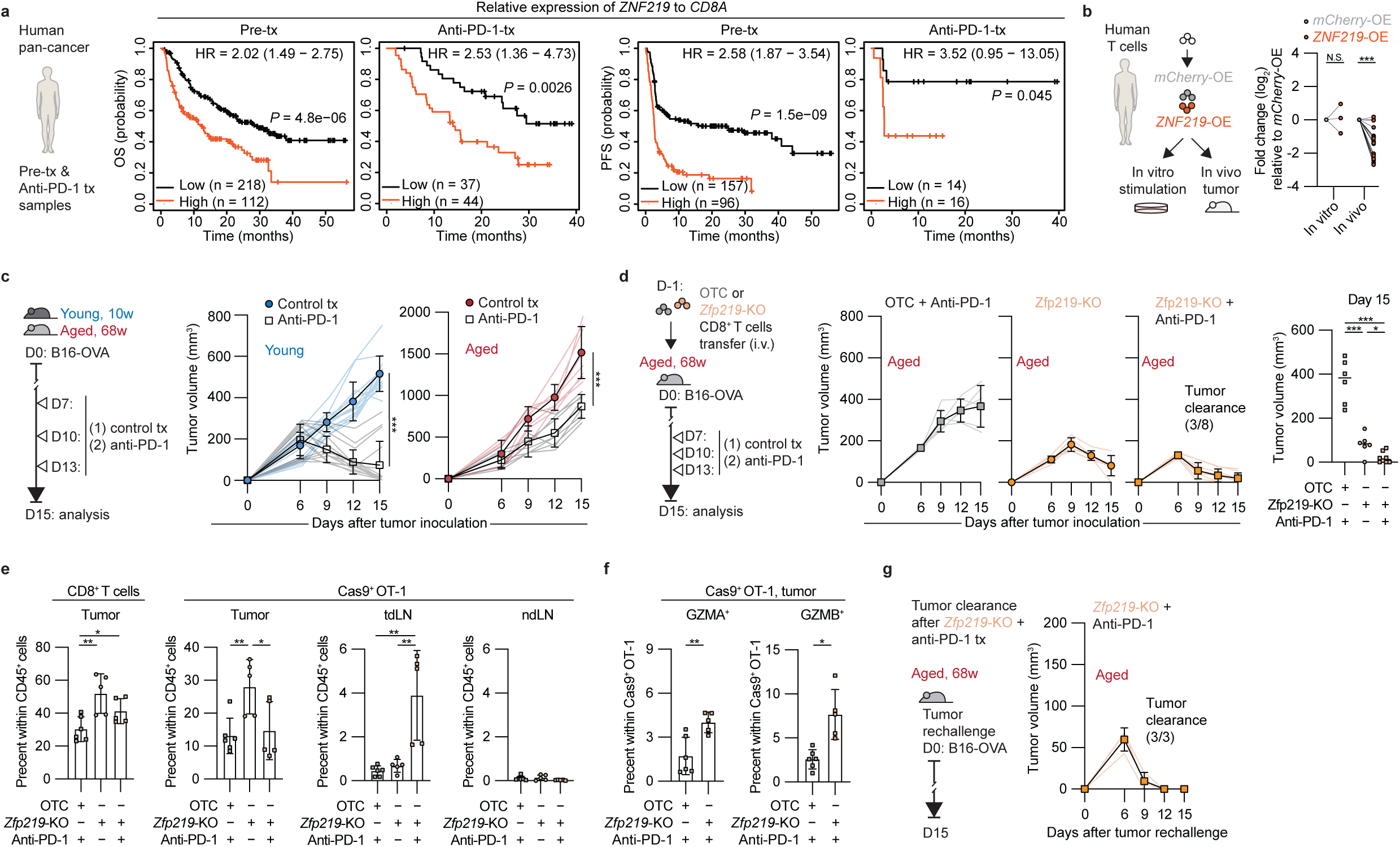
*Zfp219* KO enhances responses to immune checkpoint blockade in aged hosts. **a**, Kaplan–Meier curves illustrate overall survival (OS) and progression-free survival (PFS) related to *ZNF219* expression in intratumoral CD8^+^ T cells, based on the human pan-cancer dataset. This includes tumor biopsy samples taken before treatment (pre-tx) and samples collected during or after anti-PD-1 treatment (anti-PD-1 tx). The statistical significance was determined using the log-rank (Mantel–Cox) test. **b**, LFC of human T cells overexpressing *ZNF219* (*ZNF219*-OE) relative to *mCherry*-overexpressed (*mCherry*-OE) T cells following either 14 days of in vitro T cell stimulation (n = 3 per group) or recovery from day 14 tumors in vivo (n = 20 per group). **c**, Schema and growth curves after anti-PD-1 treatment in young (10 weeks; n = 14-15 per group) and aged (68 weeks; n = 8 per group) B16-OVA tumor-bearing mice. **d**, Schema and growth curves of aged mice (68-weeks) that were treated with *Zfp219*-KO T cells alone (n = 6), *Zfp219*-KO T cells in combination with anti-PD-1 treatment (n = 8), or OTC T cells plus anti-PD-1 treatment as a control (n =6). The final B16-OVA tumor volumes were measured on day 15. **e**, Percentage of day 15 total CD8^+^ T cells in tumors, along with OT-1 CD8^+^ T cells in tumors and tdLNs across different treatment groups (n = 5-6 per group). **f**, Percentage of day 15 GZMA^+^ or GZMB^+^ CD8^+^ T cells in tumors (n = 5-6 per group). **g**, Schema and growth curves of tumor rechallenge. One week following tumor clearance, mice that eliminated B16-OVA tumors through the combination of *Zfp219*-KO T cells and anti-PD-1 treatment were rechallenged with B16-OVA tumor cells (n = 3). Mean ± s.d. For **c-f**, significance was calculated using a two-sided Student’s t-test. Asterisks used to indicate significance correspond to the following: N.S. (not significant, *P* > 0.05), \**P* ≤ 0.05, \*\**P* ≤ 0.01, and \*\*\**P* ≤ 0.001.

The above results indicate that elevated ZNF219 expression in CD8^+^ T cells is associated with poor clinical outcomes in patients receiving anti-PD-1 therapy and intrinsically limits T cell persistence. Given that sustained CD8^+^ T cell persistence is a critical determinant of response to PD-1 blockade, we next asked whether genetic ablation of *Zfp219* (human *ZNF219*) could enhance anti-PD-1 treatment outcome in aged tumor-bearing mice. Consistent with previous studies^2,4^, anti-PD-1 treatment alone was inefficient in aged tumor-bearing mice compared with the young cohort (Fig. 5c). We then compared aged tumor-bearing mice that were treated with *Zfp219*-KO T cells alone, *Zfp219*-KO T cells in combination with anti-PD-1 treatment, or OTC T cells plus anti-PD-1 treatment as a control. Tumor control was significantly improved by the combination of *Zfp219* ablation and PD-1 blockade, with complete tumor clearance observed in 3 of 8 mice (Fig. 5d). This synergistic effect of the combination treatment was accompanied by increased intratumoral CD8⁺ T cell frequencies, robust expansion of OT-1 T cells in tdLNs, and higher proportions of GZMA⁺ and GZMB⁺ OT-1 T cells within tumors (Fig. 5e-f), indicative of enhanced cytotoxic function. Remarkably, mice that achieved complete tumor regression following the combined treatment were resistant to tumor rechallenge (Fig. 5g), demonstrating durable long-term antitumor immunity.

In summary, here we use in vivo CRISPR screens with single-cell readouts on tumor-infiltrating T cells from both young and aged mice to identify Dusp5 and Zfp219 as key suppressors of T cell persistence and effector function. We further delineate distinct mechanisms of action for these two targets, where Dusp5 limits T cell proliferation through ERK signaling, and Zfp219 restrains cytotoxic gene expression and granzyme production. Finally, we demonstrate that *Zfp219* deletion synergizes with PD-1 blockade to elicit durable antitumor immunity and prolonged tumor control in aged hosts.

## Discussion

As individuals age, T cell immunity declines, impairing effective antitumor responses through both intrinsic defects within T cells and extrinsic influences from the aged microenvironment. We previously showed that external signals in the aged TME drive CD8^+^ T cells toward an age-related dysfunction. This dysfunction differs from typical T cell exhaustion in tumors, as it is distinguished by its functional, transcriptional, and epigenetic features^1^. However, at that time, the molecular drivers behind this age-related dysfunction remained unknown. To uncover potential regulators of age-related T cell dysfunction, we performed pooled loss-of-function in vivo single-cell CRISPR screens. This approach enabled us to identify Dusp5 and Zfp219 as key regulators that restrict T cell persistence and effector function in aged hosts. Dusp5 deletion enhanced T cell proliferation via activation of ERK signaling, while Zfp219 deletion reprogrammed T cells to elevate cytotoxic granzyme expressions, leading to improved tumor control in aged tumors.

Beyond the classical protein tyrosine phosphatases (PTPs), DUSPs also play critical roles in modulating MAPK signaling through their dephosphorylation activity. The functions of DUSP family proteins have been characterized in CD4^+^ T cells^34,35^, but their roles in CD8^+^ T cell biology remain poorly defined. Here, we observed distinct roles for Dusp4 and Dusp5 in regulating CD8^+^ T cell persistence within aged tumors. Loss of *Dusp1* (also known as *MKP-1*) did not enhance CD8^+^ T cell persistence. Previous evidence suggests that *Dusp1* KO prevents NFATc1 nuclear translocation and constrains TCR signaling^36^. This signaling defect inhibits T cell activation and likely explains why *Dusp1* loss did not improve T cell persistence. By contrast, deletion of *Dusp4* or *Dusp5* enhances T cell persistence, but drives T cells toward separate differentiation trajectories. *Dusp4* KO drives CD8^+^ T cells toward the T_TAD_ state, whereas *Dusp5* KO promotes T_Eff–like_ differentiation. This divergent result suggests potentially opposing regulatory roles for Dusp4 and Dusp5 in shaping T cell fate through distinct MAPK signals. Moreover, Dusp5, a nuclear phosphatase, may function as a broadly applicable, druggable regulator of T cell reprogramming by selectively constraining nuclear MAPK activity, thereby shaping downstream transcriptional programs that govern T cell fate and persistence.

Our study identifies Zfp219 as a previously unrecognized regulator of CD8^+^ T cell function. The zinc finger protein (Zfp; also known as human ZNF) family exhibits distinct structures and functions, including RNA-binding and DNA-binding transcriptional regulators. Zfp219 (ZNF219) in particular has been reported primarily as a DNA-binding transcriptional repressor^31,32^. Here, we observed that Zfp219 expression is markedly increased in tumor-infiltrating CD8^+^ T cells from both human cancer patients and aged tumor-bearing mice, suggesting that the aged environment promotes its upregulation. This probably explains why the better tumor control and survival benefit upon *Zfp219* KO are observed only in aged, but not young, hosts. Mechanistically, our results suggest that Zfp219 acts as a repressor of key cytotoxic effector genes, including *Gzma* and *Gzmb*. Loss of Zfp219 in T cells restores granzyme production and enhances cytotoxic function, thereby rejuvenating antitumor immunity in aged settings. These findings position Zfp219 as a potential therapeutic target to reinvigorate CD8^+^ T cell cytotoxicity in aged tumors and highlight the contribution of zinc finger transcriptional regulators to immune aging and cancer therapy. More broadly, our findings position zinc finger transcriptional regulators, such as Zfp219 (ZNF219), as potential targets for restoring T cell function in immune aging associated diseases.

One limitation of our study is the relatively small size of the CRISPR library, which comprised 60 target genes and two control genes. These genes were selected based on previously identified transcriptional differences in intratumoral CD8^+^ T cells between young and aged tumors. There are two major bottlenecks associated with in vivo single-cell CRISPR screening that constrain scalability to the genome-wide level. The first is the limited recovery of T cells from tumors. To maximize T cell engraftment and T cell recovery from tumors while avoiding tumor clearance, we optimized the highest possible dose of T cells per mouse. The second restriction arises from the high cost and limited throughput of the Perturb-seq platform. Together, these considerations make it challenging to scale in vivo single-cell CRISPR screens to a genome-wide level, providing the rationale for downsizing our library to a more targeted set of relevant genes in the current study.

In conclusion, our study identifies Dusp5 and Zfp219 as key suppressors of CD8^+^ T cell function within aged tumors. Deletion of either gene increases T cell persistence and promotes differentiation toward an effector state. Notably, loss of *Dusp5* enhances T cell proliferation by activating ERK signaling, while loss of *Zfp219* reprograms T cells to unleash cytotoxic granzyme expression. Together, these findings identify Dusp5 and Zfp219 as potential targets to improve T cell-mediated tumor control in aged hosts and highlight therapeutic opportunities in combination with ICB treatment to boost antitumor immunity in older individuals.

## Methods

### Study design

This study was designed to identify genes that, when deleted, enhance the CD8^+^ T cell persistence and function in targeting tumors, including melanoma and lung cancer, in aged tumor-bearing mice. To validate the targets identified in the loss-of-function in vivo single-cell CRISPR screen, functional assays were conducted to confirm the underlying mechanisms. All animal procedures were performed in accordance with protocols approved by the Massachusetts General Hospital (MGH) Institutional Animal Care and Use Committee (IACUC). Animals were euthanized in accordance with the experimental protocol or when they reached pre-specified endpoints as defined by the IACUC. The maximum permissible tumor size was a diameter of 20 mm, which was not exceeded. Mice used for in vivo experiments were randomized prior to T cell transfer and tumor inoculation.

### Mice

All in vivo experiments were performed with female mice. Female C57BL/6J mice (strain 000664), aged C57BL/6J mice (stock 000664), CD45.1 congenic mice (B6.SJL-Ptprc^a^Pepc^b^/BoyJ, strain 002014), OT-1 transgenic mice (C57BL/6-Tg(TcraTcrb) 1100Mjb/J, strain 003831), Cas9 mice (B6J.129(Cg)-Gt(ROSA)26Sor^tm1.1(CAG–cas9*,–EGFP)Fezh^/J, strain 026179), and NSG mice (NOD.Cg-Prkdc^scid^ Il2rg^tm1Wjl^/SzJ, strain 005557) were purchased from The Jackson Laboratory. Cas9 OT-1 CD45.1 mice were cross-bred and routinely maintained. All breeding mice and subsequent litters were genotyped at Transnetyx. All experimental mice were housed under specific pathogen-free conditions, used in accordance with ethical regulations, and approved by the MGH IACUC.

### Cell lines and tumor inoculation

B16-OVA and LLC-OVA cells modified to express OVA were gifts from B. Miller (University of North Carolina, Chapel Hill) and R. Manguso (Broad Institute), respectively. Lenti-X HEK293 cells were purchased from TaKaRa Bio (632180). A375 melanoma cells were purchased from the American Type Culture Collection (ATCC). A375 melanoma cells expressing luciferase and membrane-tethered anti-CD3 scFv and human CD80 were prepared as the previous study^37^. Cells were cultured in DMEM (Thermo Fisher Scientific, 11995073) supplemented with 10% fetal bovine serum (FBS), 10 mM HEPES (Thermo Fisher Scientific, 15630080), 1% penicillin/streptomycin (Thermo Fisher Scientific, 15140122), and maintained at 37°C with 5% CO2. Cells were routinely tested for Mycoplasma contamination (and were negative) using a LookOut Mycoplasma PCR detection kit (Sigma), and were also authenticated by STR profiling in a three-year cycle. Mice were subcutaneously injected with 1 × 10^6^ B16-OVA cells or 1 × 10^6^ LLC-OVA cells in 1× HBSS (Life Technologies). Tumors were measured every 3 days starting at day 6 after tumor injection. Tumor volume (mm^3^) was calculated using the following formula: *V* = *L* × *W* × *W*/2, where *L* and *W* denote length and width, respectively.

### In vivo transfer with anti-PD-1 treatment

Naive young *Dusp5*-KO, *Zfp219*-KO, and OTC Cas9^+^ OT-1 CD45.1^+^ CD8^+^ T cells were sorted on a FACSAria Fusion (BD Biosciences) on day 3 after virus transduction. 5,000 cells of each cell type were intravenously injected into female CD45.2^+^ young and aged C57BL/6J mice one day before OVA-tumor inoculation. OVA-tumor cells were subcutaneously implanted with 1 × 10^6^ cells. Tumor-bearing mice were treated with 100 μg anti-PD-1 (clone 29F.1A12, BioXCell) or rat IgG2a isotype control antibody (clone 2A3, BioXCell) on days 7, 10, and 13. Tumors were harvested on day 15 for analysis.

### Lentiviral vector and pooled gRNA cloning

The lentiviral vector (pRDA_526) allows continuous sgRNA expression and includes Thy1.1 as a transduction marker, which is available on Addgene (ID: 184859)^16^. An sgRNA cassette with the human U6 promoter was modified for T cell screening by incorporating a fixed tracrRNA following additional gRNA capture sequencing, and adding BsmBI sites compatible with Golden Gate cloning for easy insertion of variable sgRNAs. We replaced Thy1.1 with the fluorophore mAmetrine (also called Vex).

The gRNA oligonucleotides used in the in vivo CRISPR screens were designed using the Broad CRISPR platform - [CRISPick](https://portals.broadinstitute.org/gppx/crispick/public). We designed 6 gRNAs per gene and included gene-targeting sequences for 60 genes encoding nuclear proteins, 2 control genes (*Pdcd1*, *Ptpn2*), 20 non-targeting control (NTC) sequences, and 20 one-targeting control (OTC, intergenic control) sequences (as Extended Data Fig. 1e). All sgRNA sequences in the pool are listed in Supplementary Table 2. The UMI oligonucleotides were created as all possible permutations of 6 bp sequences and were added after the gRNA oligonucleotides to verify the diversity of sgRNA after pool gRNA cloning.

pRDA_526 was digested with BsmBI-v2 (NEB, R0739S). gRNA libraries, including UMI sites, were prepared using IDT oPools service and cloned into the digested pRDA_526 via NEB Golden Gate Assembly. Ligation products were purified using the MiniElute PCR purification kit (QIAGEN, 28004). These products were then electroporated into Endura Competent Cells (Biosearch Technologies, 60242-2), which were seeded into 500 mL Luria Broth (with ampicillin) and incubated overnight at 30°C. Plasmid libraries were extracted using the ZymoPURE II Plasmid Maxiprep Kit (ZymoPure, D4203), quantified with Qubit™ dsDNA Quantification Assay Kits (Thermo Fisher Scientific, Q32851), and sequenced to ensure even gRNA representation on an Illumina MiSeq with custom primers (single-end 40-8).

### Lentiviral production and transduction

Lenti-X HEK293 cells were transfected using lipofectamine 3000 (Thermo Fisher Scientific, L3000015) with the packaging plasmids psPAX2 (Addgene, 12260), pMD2.G (Addgene, 12259), pCag-Eco (Addgene, 35617), and pRDA_526 plasmid containing the gRNA libraries. The above plasmids were used at a 2:1:1:6 mass ratio to produce the pseudotyped lentivirus. Lentivirus-containing supernatant was collected 54-72 hours after transfection, spun at 500 x g for 5 minutes to remove cells, and concentrated by Lenti-X Concentrator (TaKaRa Bio, 631232) overnight. Non-tissue culture-treated 24-well plate was coated with RetroNectin (Takara Bio, T100B) at 15 μg/cm^2^ at 4°C overnight. The next day, the RetroNectin was taken out from the 24-well plate, 500 µl of 2x concentrated lentivirus was added to the RetroNectin-coated plate, and it was centrifuged at 2,000 g for 2 hours at 32°C.

Naive young 10-week-old Cas9^+^ OT-1 CD8^+^ T cells were isolated from spleens using the CD8a T cell Isolation Kit (Miltenyi BioTec, 130-104-075). They were then plated on virus-coated plates with 1 μg/ml LentiBOOST-P (Mayflower Bioscience, SB-P-LV-101-12) in complete RPMI media, which includes RPMI-1640 (Thermo Fisher Scientific, 11875119), 10% FBS, 1% penicillin/streptomycin, 10 mM HEPES, 55 μM 2-mercaptoethanol (Thermo Fisher Scientific, 21985023), and 10 ng/ml recombinant murine IL-7 (PeproTech, 217-17). Cells were cultured for 3 days to enable T cell expression of mAmetrine while maintaining cell viability and naïve cell status. Fresh media and cytokine were exchanged on day 2 after transduction. mAmetrine^+^ Cas9^+^ OT-1 CD8^+^ T cells were sorted on a FACSAria Fusion (BD Biosciences) on day 3 after transduction. Genomic DNA was isolated from 5 × 10^4^ CD8^+^ T cells using the QIAamp DNAMicro Kit (Qiagen, 56304), as per the manufacturer’s protocol. After PCR, next-generation sequencing was performed (complete amplicon sequencing) by the MGH DNA Core.

### In vivo loss-of-function CRISPR screen

Young 10-week-old and aged 68-week-old mice received intravenous injections of sorted mAmetrine^+^ Cas9^+^ OT-1 CD8^+^ T cells (as described above) one day before B16-OVA tumor inoculation. On day 15 after tumor cell injection, tumors were dissected from the surrounding fascia, mechanically minced, and treated with collagenase P (2 mg ml^−1^; Sigma) and DNase I (50 μg ml^−1^; Sigma) for 15 min at 37 °C. After tumor samples were filtered with a 70-μm filter, tumor-infiltrating lymphocytes underwent red blood cell lysis (RBC lysis buffer, Biolegend), and were further enriched by CD8-positive MACS selection (Miltenyi Biotec, 130-117-044). tdLNs were dissected, mechanically minced, and filtered into single-cell suspensions. When necessary, CD8^+^ T cells were sorted on a FACSAria Fusion (BD Biosciences) for downstream Perturb-seq and/or ATAC-seq analyses.

### CD8^+^ T cell isolation from tissues

On the indicated days after tumor cell injection, tumors were dissected from the surrounding fascia, mechanically minced, and treated with collagenase P (2 mg ml^−1^; Sigma) and DNase I (50 μg ml^−1^; Sigma) for 15 min at 37 °C. After tumor samples were filtered with a 70-μm filter, tumor-infiltrating lymphocytes were performed red blood cell lysis (RBC lysis buffer, Biolegend), and were further enriched by CD8-positive MACS selection (Miltenyi, 130-117-044). tdLNs were dissected, mechanically minced, and filtered into single-cell suspensions. When necessary, CD8^+^ T cells were sorted on a FACSAria Fusion (BD Biosciences) for downstream Perturb-seq and/or ATAC-seq analyses.

### Flow cytometry and antibodies

To investigate phenotypic and functional changes by flow cytometry, cells were blocked with anti-mouse CD16/CD32 (Biolegend) and surface stained with the indicated antibodies (Supplementary Table 1) for 30 min at 4 °C. NIR Live/Dead (Thermo Fisher Scientific, L34975) dye, used to exclude dead cells from downstream flow cytometry analysis, was added in parallel with surface antibodies. To investigate intracellular markers, cells were fixed and permeabilized using a BD Cytofix/Cytoperm Fixation/Permeabilization Kit (BD Biosciences, 554714), as per the manufacturer’s instructions. Cells were blocked again with 5% mouse and rat serum plus anti-mouse CD16/CD32 and stained with intracellular antibodies for 30 min at 4 °C.

For BrdU experiments, mice were injected with 1 mg of BrdU (BD Pharmigen, 552598) intraperitoneally 16h before tissue collection. BrdU staining was performed after cell-surface staining, as per the manufacturer’s protocol.

For PhosFlow experiments, live tumor-infiltrating CD8⁺ T cells were collected from tumors on day 15. Cells were stained with surface marker antibodies with NIR Live/Dead 30 min on ice, fixed with 2% paraformaldehyde for 10 min at room temperature. For intracellular phospho-protein staining, fixed cells were permeabilized with ice-cold BD PhosFlow Perm Buffer III (BD Biosciences, 558050) for 60 min, then stained with phospho-specific antibodies in BD Perm/Wash buffer (BD Biosciences, 554723) for 1 h at room temperature, as per the manufacturer’s protocol.

A BD Fortessa X-20 (BD Biosciences) flow cytometer and FACSDiva software (v.8.0.1, BD Pharmingen) were used to collect data, and analyses were performed using FlowJo software (v.10.10.0, BD). Single-color compensation controls and fluorescence minus one thresholds were used to set flow gate margins.

### Perturb-seq

Live tumor-infiltrating mAmetrine^+^ Cas9^+^ OT-1 CD8^+^ T cells were sorted from tumors and tdLNs on day 15 after tumor inoculation by a BD FACSAria Fusion Sorter (BD). Sorted cells were pooled into groups (2 biological replicates per sample and 4 samples per age group). Cells were counted and loaded onto the Chromium X (10x Genomics) for a target recovery of 10,000 single cells per run. Samples were then processed using the 10x Chromium Next GEM Single Cell 5’ Reagent Kit v2 (Dual Index, CG000511 Rev B) with Feature Barcode technology for CRISPR screening. Library fragment sizes were assessed with TapeStation HS1000. Sequencing was performed on an Illumina NovaSeq X instrument.

### Perturb-seq analysis

The CellRanger analysis pipeline (v.7.0.1) was used to demultiplex samples, process barcodes, align reads, filter reads, count unique molecular identifiers, and aggregate sequencing runs. Transcripts were aligned to the refdata-gex-mm10-2020-A, a mouse transcriptome (CellRanger version 2020-A). sgRNA reads were aligned to the library using the pattern TCAAAC(BC)CCCCAT. Downstream analyses were performed in R (v.4.5.1) using the Seurat package (v.4.4.0). In downstream analysis, cells with fewer than 200 unique genes, greater than 5,000 unique genes, greater than 10% mitochondrial reads, and more than 2 unique sgRNA were removed, yielding, on average, an expression matrix of 49,350 cells by 32,285 genes.

Gene expression measurements were normalized by total expression per cell, multiplied by a scaling factor of 10,000, and transformed using log (1*p*). The top 3,000 most variable features were selected using the ‘vst’ method in Seurat’s FindVariableFeatures for PCA analysis. The top PCs were calculated and determined to be significant (P < 0.01) using the ‘elbow plot’ method. A UMAP was calculated from 15-20 PCs using default parameters. Unsupervised clustering was performed using the original Louvain algorithm. Multiple clustering resolutions were examined, and an optimal resolution of between 0.4 and 0.6 for clustering was selected for further analysis. All differential expression analysis was performed using DESeq2 (PyDESeq2v.0.4.8), and volcano plots were made using EnhancedVolcano (v.1.18.0) and ggplot2 (v.3.4.2). VISION (v3.0.1) was used to create an age-associated CD8^+^ T cell dysfunction gene signature library, and this library was used to predict the similarity between age-associated CD8^+^ T cell clusters in tumors and tdLNs. To visualize the rank of top-performing guides for each condition comparison, the values were plotted in a frequency histogram by the sgRankView function in MAGeCKFlute (v2.12.0). Ranked lists of differentially expressed sgRNAs were created from log_2_ fold change (log_2_FC) values calculated with DESeq2 (v1.48.2).

### scRNA-seq analysis

The published mouse scRNA-seq dataset was collected from our previous GSE255373. The published human pan-cancer scRNA-seq datasets were collected from E-MTAB-8017, GSE115978, GSE120575, and GSE179994. A summary of human cancer types and age information can be found in Supplementary Table 3. Downstream analyses were performed in R (v.4.5.1) using the Seurat package (v.4.4.0). In downstream analysis, cells with fewer than 200 unique genes, greater than 5,900 unique genes and greater than 10% mitochondrial reads were removed and normalized by sctransform (v.0.3.5), and the CD8^+^ T cell cluster was purified. To perform differential gene expression analysis at the individual-sample level, a pseudobulk approach was used. For each sample, raw UMI counts from single cells were aggregated by summing across all cells from the same individual, yielding a sample-specific count matrix. The aggregated counts were then normalized to reads per million (RPM) to account for differences in sequencing depth. Normalized expression values were compared between experimental and control groups by calculating the log_2_FC for each gene.

### ATAC-seq library preparation and analysis

Live tumor-infiltrating mAmetrine^+^ Cas9^+^ OT-1 CD8^+^ T cells from tumors on day 15 after tumor inoculation were sorted and pooled into two or three independent biological replicates, which ranged from 10,000 to 30,000 cells per subset combination (T_Prog_: SLAMF6^+^TIM-3^−^; T_Tran_/T_Eff-like_: SLAMF6^+^TIM-3^+^; T_Term_: SLAMF6^−^TIM-3^+^; T_TAD_: SLAMF6^−^TIM-3^−^). As previously described, pelleted cells were incubated in 5–50 μl of reaction mix containing 2×TD, Tn5 enzyme, and 2% digitonin in nuclease-free water at 37 °C for 30 min with agitation at 1,000 rpm. A MinElute Reaction Cleanup kit (Qiagen) was used for DNA purification. Following PCR, bead cleanup was performed using AMPure XP beads (Beckman Coulter/Agencourt), library quality was verified by TapeStation analysis, and library quantification was confirmed using the KAPA Library Quantification kit (Illumina Platforms; Roche). Sequencing was performed on a NextSeq1000/2000 sequencer using paired-end 50-base pair reads.

Raw fastq files were downloaded from Illumina BaseSpace and merged across lanes. FastQC was run before and after trimming. Trimmomatic (v.0.36) was used for quality trimming and primer removal in raw fastq files with the following parameters: LEADING: 15, TRAILING:15, SLIDINGWINDOW: 4:15, MINLEN: 30, and CROP: 35. Trimmed reads were aligned with Bowtie2 (v.2.2.9) to mm10, with a maximum insert size of 1,000 bases. Aligned bam files were sorted, and duplicates were marked (Picard v.2.8.0). Reads that mapped to the blacklist region were removed. MACS (v.2.1.1) was used to perform peak calling on merged BAM files (Samtools v.1.3.1) from biological replicates using a q-value threshold of 0.001. TDF files were generated using igvtools (v.2.3.98) for visualization in the Integrative Genomics Viewer (v.2.13.1, Broad Institute). Consensus peaks from all samples were merged to create a single peak universe. A raw counts matrix was generated for each biological replicate by quantifying the number of cut sites within each peak region. The ataqv package was used for data quality control, including transcription start site enrichment. The Combat-seq method was used to batch correct samples sequenced at different time points by Surrogate Variable Analysis (sva, v.3.48.0), and peaks with fewer than 25reads across all samples were removed. DESeq2 (v.1.40.2) was used for count matrix normalization and differential accessibility analysis across pairwise comparisons of all conditions. R (v.4.3.1) was used to perform PCA. Gene-to-peak associations were determined using the rGREAT package (v.3.2.1) with default settings. rGREAT was also used to determine the gene set enrichment, with default settings and hypergeometric and binomial tests to measure significance.

### Multiplex immunofluorescence

FFPE tissue sections from B16-OVA melanoma were stained on a Leica Bond RX automated stainer. Antigen retrieval was performed with Bond ER2 solution (Tris-EDTA buffer, pH 9.0) for 40 minutes. The multiplex immunofluorescence (IF) panel used the Opal 6-Plex Anti-Rabbit Detection Kit (Akoya NEL871001KT) and rabbit anti-mouse CD8α monoclonal (clone D4W2Z, Cell Signaling #98941) with Opal 780. The primary CD8α antibody was diluted 1:200, and Opal 780 fluorophore was diluted 1:25. DAPI was used for nuclear staining. Whole-slide images were captured on the PhenoImager HT multispectral slide scanner (Akoya Biosciences) at 20x magnification with a resolution of 0.50 µm per pixel. Imaging settings were optimized for each slide to avoid pixel saturation or underexposure. Staining was quantified at the cellular level within the annotated regions using the HALO HighPlex FL module v4.1.3 and HALO AI module v3.6.1434. Nuclear detection and cell segmentation were performed with the default AI nuclear segmentation algorithm based on the DAPI channel. For each cell, positivity thresholds for each antibody marker in the cytoplasmic and nuclear compartments were determined based on signal intensity and area coverage.

### Human survival analysis

Gene expression data from all eligible human cancer datasets were combined, normalized, and scaled to 1000, as previously described^33^. Tumor samples obtained before therapy were termed “pre-treatment (pre-tx),” and those collected during or after anti-PD-1 therapy were termed “anti-PD-1 treatment (anti-PD-1 tx).” Progression-free survival (PFS) was calculated from the date of treatment until evidence of progression, as determined by the clinician, leading to treatment discontinuation/switch or death, whichever occurred first. Survival time was not used if the patient had no event, and follow-up time was censored before 12 months. Overall survival was calculated from the date of treatment until the date of death or censure.

### Human T cell studies

Healthy donor primary T cells were isolated from leukopaks purchased from STEMCELL Technologies and obtained with consent from human donors, using the EasySep Human T cell isolation kit (STEMCELL Technologies, 17951). Isolated T cells were activated with Dynabeads Human T-Activator CD3/CD28 for T Cell Expansion and Activation (Thermo Fisher Scientific, 11132D) at a 1:1 ratio of cells to beads for 48 hours. T cells were transduced with lentivirus after 24 hours of activation and expanded in recombinant IL-2 (100 IU) for 12 days before freezing. To evaluate the impact of ZNF219 overexpression, T cells expressing lentiviral vectors encoding mCherry (control) or ZNF219 were thawed and injected intravenously into NSG mice with established A375 anti-CD3/CD80 tumors. A375 anti-CD3/CD80 cells were engineered to express membrane-bound OKT3 scFV and CD80, enabling universal human T cell engraftment. NSG mice were subcutaneously injected with 2 × 106 cells in 50% matrigel. 5 × 106 T cells were injected 14 days after tumor implantation and were reisolated using the EasySep Release Human CD45 Positive Selection Kit (STEMCELL Technologies, 100-0105) 14 days later. Alternatively, T cells were restimulated with Immunocult Human CD3/CD28 T Cell Activator (STEMCELL Technologies, 10971) and expanded in IL-2 for 14 days.

### Statistical analysis

Sample sizes of five to ten animals per experimental arm per experimental replicate were chosen to ensure adequate power. Data collection and analysis were not performed blind to the conditions of the experiments. We performed statistical analyses with GraphPad Prism 10 (v.10.6.1) or R. Unless otherwise stated, data are presented as mean ± s.d., and a two-tailed Student’s t-test was used. All tests were two sided unless otherwise specified. Significance was considered for *P value of* < 0.05 (\**P* < 0.05, \*\**P* < 0.01, and \*\*\**P* < 0.001).

## Data availability

Sequence data (Perturb-seq and ATAC-seq) have been deposited in the Gene Expression Omnibus under accession codes GSE306058 and GSE310273. All human scRNA-seq datasets analyzed in this study have been previously published and are publicly available (E-MTAB-8017, GSE115978, GSE120575, and GSE179994). Source data are provided with this paper.

## Code availability

No custom code beyond adaptation of existing software packages was used in this study. All code will be provided upon reasonable request by the corresponding author.

## Acknowledgements

We thank all members of the Sen laboratory, as well as members of the Krantz Family Center for Cancer Research, for thoughtful scientific discussions. We thank the MGH Flow Cytometry Core (especially, Meaghan A. Cruise), the MGH Animal Research Facility, the Broad Clinical Labs Sequencing Service, the MGH Pathology Core, and the Biomarker Discovery Lab for their assistance. D.R.S. was supported by funding from the Melanoma Research Alliance, the V Foundation for Cancer Research, and NIH DP2AI176139. A.C.Y.C. was supported by funding from NIH T32CA921643 and 1K99CA296769. K.Y.J is supported by National Science Foundation Graduate Research Fellowship under Grant No. (DGE 2140743).

## Author contributions

CRISPR screen conception and experimental design: A.C.Y.C., K.Y.J., C.Y., M.Z., M.A.S., N.H., and D.R.S. Performed CRISPR screen experiments: A.C.Y.C., K.J., and C.Y. Target validation and mechanistic experiments: A.C.Y.C., K.Y.J., and N.H.K. Critical reagents and advice provided by K.B.Y., R.T.M., T.R.M., M.A.S., and N.H. Animal handling: A.C.Y.C., K.Y.J., and D.M. Sample processing and preparation: A.C.Y.C., K.Y.J., D.Z., K.T., K.H.X., E.M.S., and L.T.N. Analysis and interpretation of data: A.C.Y.C., K.Y.J., K.H.X., E.M.S., and L.T.N., and D.R.S. Computational analyses: A.C.Y.C., K.Y.J., K.B., and T.J.C-L. Manuscript writing and revision: A.C.Y.C., K.Y.J., K.B.Y., R.T.M., N.H., and D.R.S.

## Corresponding author

Correspondence to Alex C. Y. Chen and Debattama R. Sen

---

**Extended Data Fig. 1.**
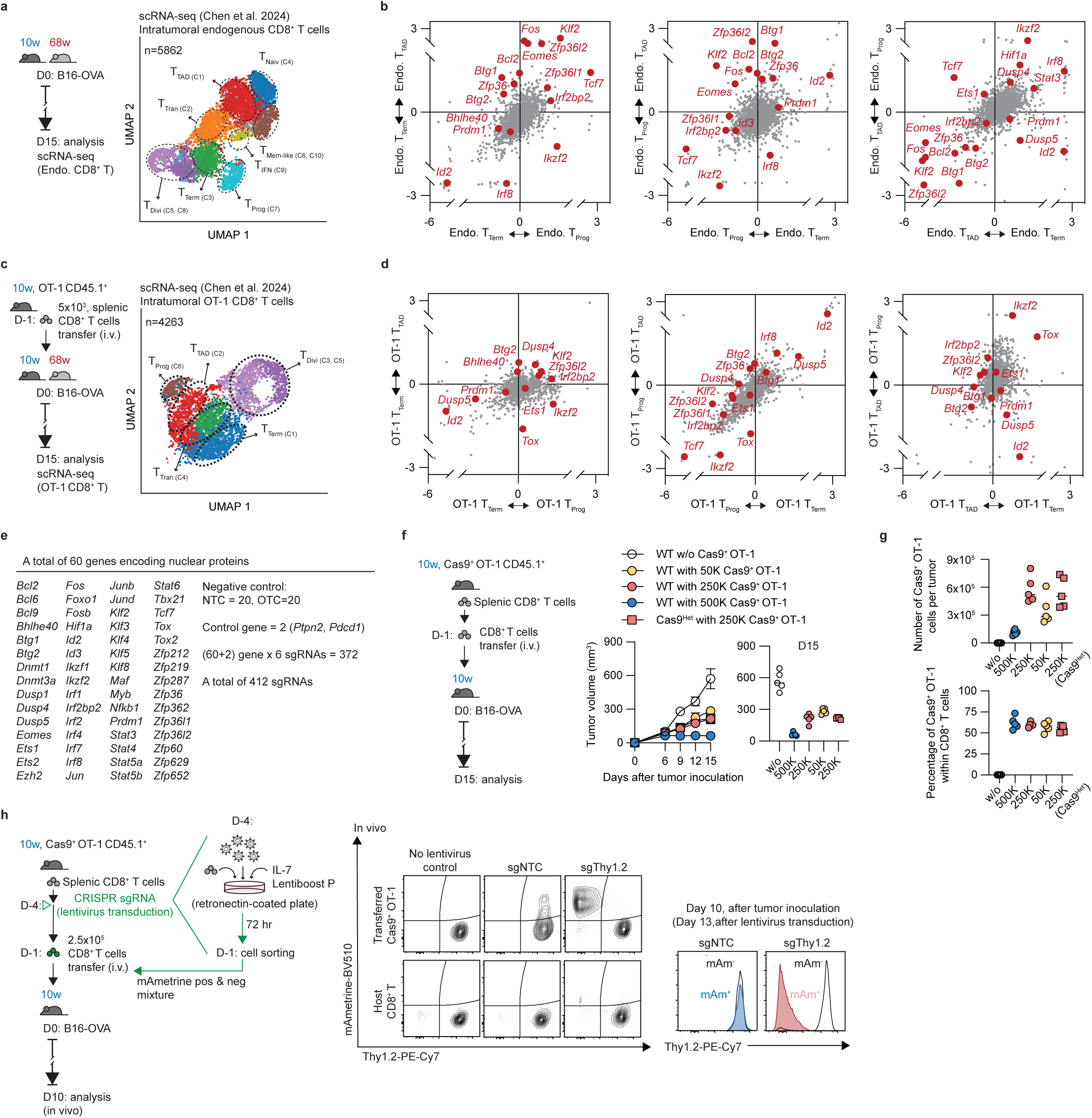
Pinpointing potential transcriptional programs of CD8^+^ T cells for in vivo scCRISPR screens. **a**, Schema of the experimental design for the B16-OVA melanoma model used to assess tumor growth in young (10 weeks; blue) and aged (68 weeks; red) mice; D0, day 0; D15, day 15. scRNA-seq UMAP projection of 5862 endogenous tumor-infiltrating CD8^+^ T cells. **b**, Log2 fold change (LFC) comparison for differentially expressed gene-encoded nuclear proteins between T cell subsets. **c**, Schema of the experimental design for B16-OVA tumors in young (10 weeks; blue) and aged (68 weeks; red) mice that received young splenic OT-1 CD8^+^ T cells. scRNA-seq UMAP projection of 4263 tumor-infiltrating OT-1 CD8^+^ T cells. **d**, LFC comparison for transcriptional regulators between OT-1 T cell subsets. **e**, A CRISPR library was designed to target 60 nuclear proteins, including *Pdcd1* and *Ptpn2* as positive controls for T cell activation. It also includes 20 non-targeting controls (NTCs), 20 intergenic one-target controls (OTCs), and 6 unique sgRNAs for each gene target, resulting in a total of 412 single-guide RNAs (sgRNAs). **f-g**, Schematic of optimizing the transfer OT-1 T cell number that could be recovered for in vivo screening. Tumor growth kinetics were monitored. **h**, Schematic illustrating gene deletion in naive T cells via an in vivo CRISPR screen platform. Includes flow cytometry plots showing Thy1.2 expression in CD8^+^ T cells from a representative non-targeting control (NTC) sgRNA or Thy1.2 sgRNA.

**Extended Data Fig. 2.**
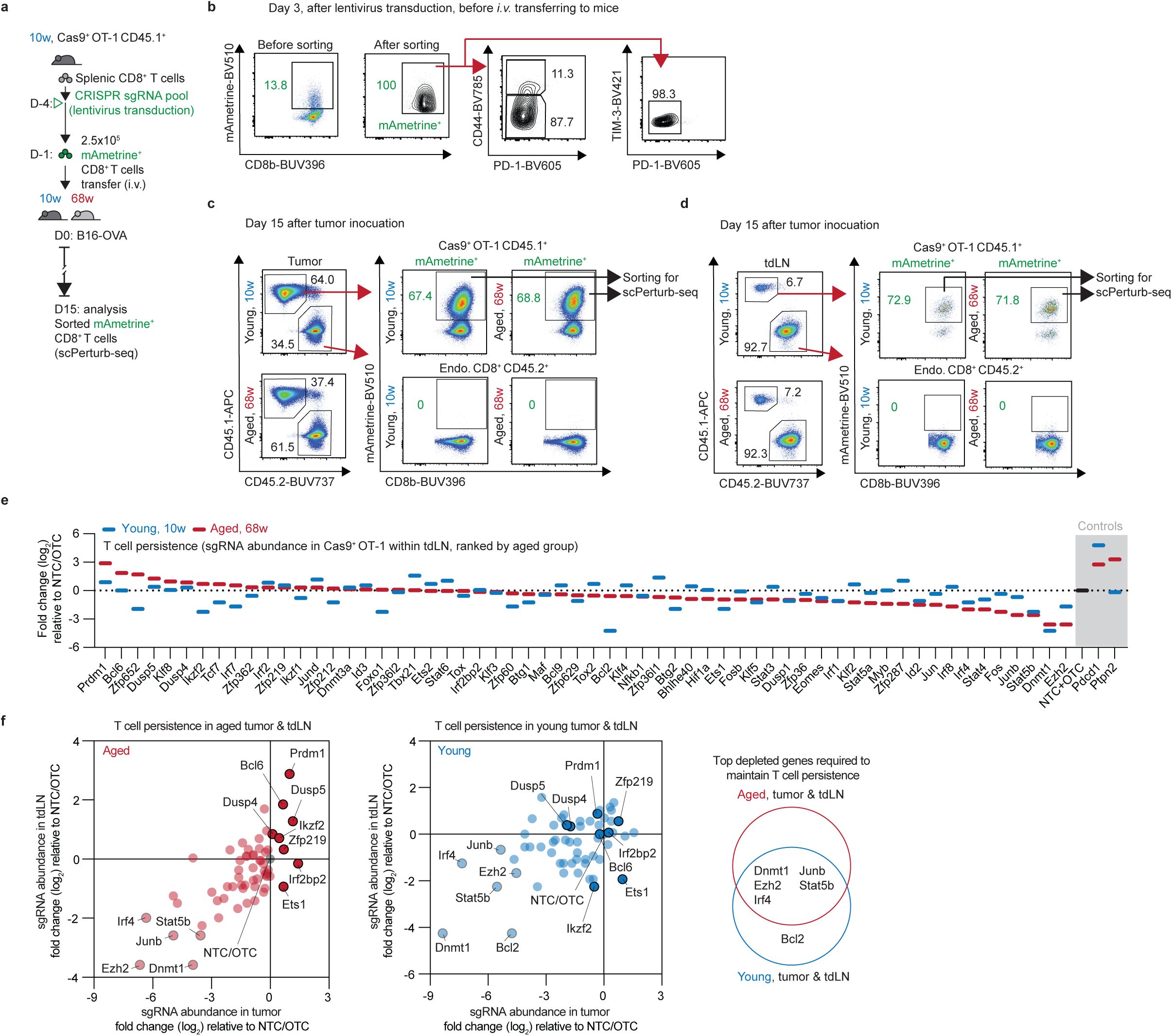
In vivo scCRISPR screens in Cas9^+^ OT-1 CD8^+^ T cells, related to Figure 1. **a**, Schematic for in vivo screening. **b**, Representative flow cytometry plots of transduced naïve mAmetrine^+^ Cas9^+^ OT-1 CD45.1^+^ CD8^+^ T cells after sorting and before intravenous (i.v.) transfer to mice. **c-d**, Representative flow cytometry plots of transduced mAmetrine^+^ Cas9^+^ OT-1 CD45.1^+^ CD8^+^ T cells after sorted from tumors or tdLNs. **e**, Log2 fold change (LFC) comparison for enriched or depleted sgRNAs of each gene, relative to NTC/OTC. Targeting genes in aged tdLNs is shown in red, and in young tdLNs in blue. **f**, LFC comparison for each targeting gene enriched in both tumors and tdLNs, relative to NTC/OTC; aged in red and young in blue.

**Extended Data Fig. 3.**
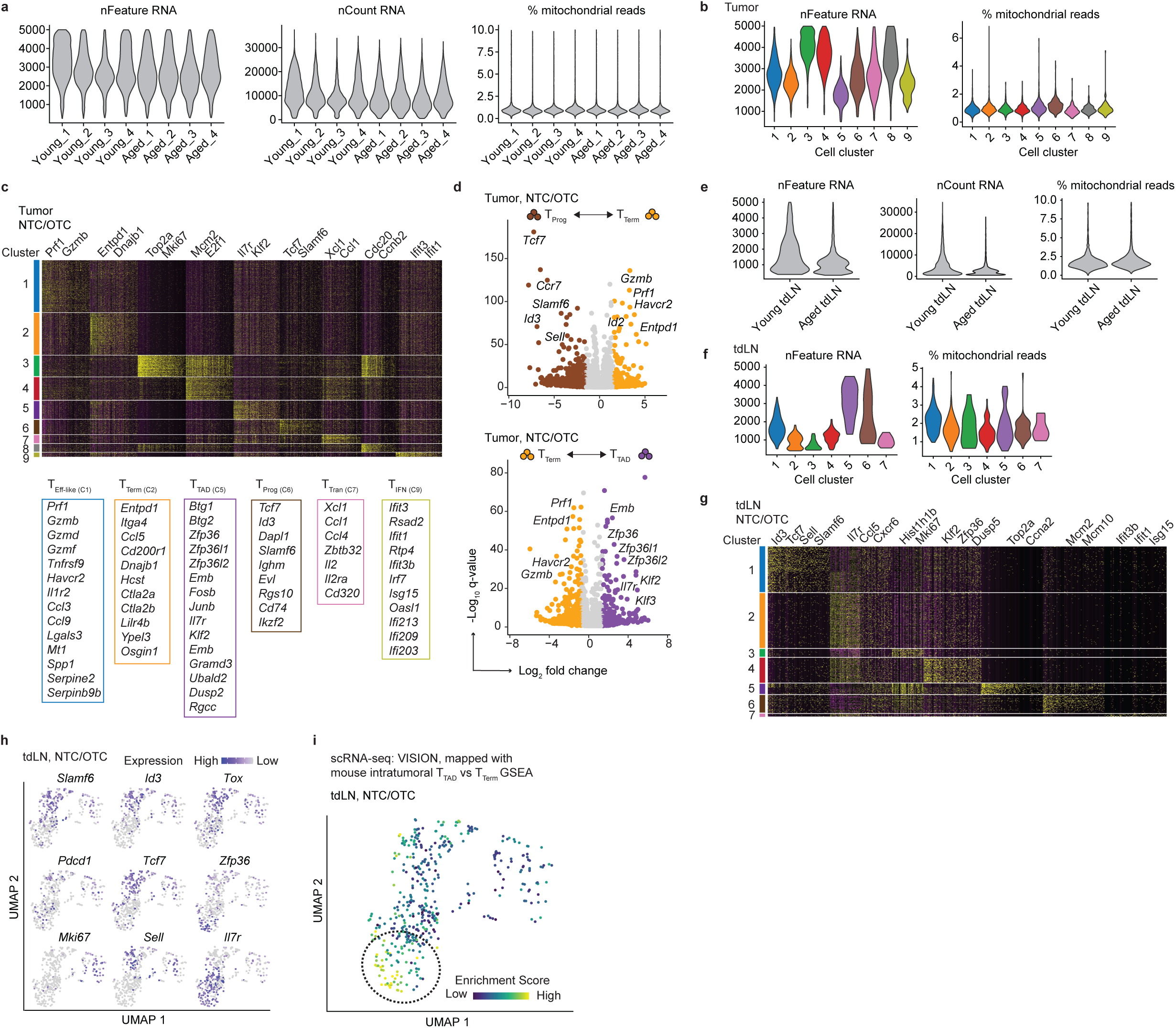
Phenotyping of Cas9^+^ OT-1 CD8^+^ T cells in young vs. aged tumors and tdLNs, related to Figures 1-3. **a-b**, Violin plots of scRNA-seq data of OT-1 CD8^+^ T cells from tumors display RNA feature counts (nFeature RNA; left), RNA read counts (nCount RNA; middle), and the percentage of mitochondrial gene reads per cell (% mitochondrial reads; right) across different groups and clusters. **c**, The heatmap shows the top cluster-defining genes in each OT-1 CD8^+^ T cell cluster/subset from tumors, as identified in Fig 2a. **d**, Volcano plot of differentially expressed genes between transferred OT-1 subsets in tumors: progenitor exhausted (T_Prog_; brown) and terminally exhausted (T_Term_; orange), as well as between T_Term_ (orange) and tumor-infiltrating age-associated dysfunctional (T_TAD_; purple) OT-1 CD8^+^ T cells. **e-f**, Violin plots of scRNA-seq data of OT-1 CD8^+^ T cells from tdLNs display RNA feature counts (nFeature RNA; left), RNA read counts (nCount RNA; middle), and the percentage of mitochondrial gene reads per cell (% mitochondrial reads; right) across different groups and clusters. **g**, The heatmap shows the top cluster-defining genes in each OT-1 CD8^+^ T cell cluster/subset from tdLNs, as identified in Fig 3i. (**h**) Expression of indicated genes in OT-1 T cells from tdLN (grey: low, purple: high). (**i**) UMAP projection showing the enrichment of mouse T_TAD_ cell signatures within mouse tdLN T cell subsets. Dark blue indicates low enrichment, while yellow-green indicates high enrichment.

**Extended Data Fig. 4.**
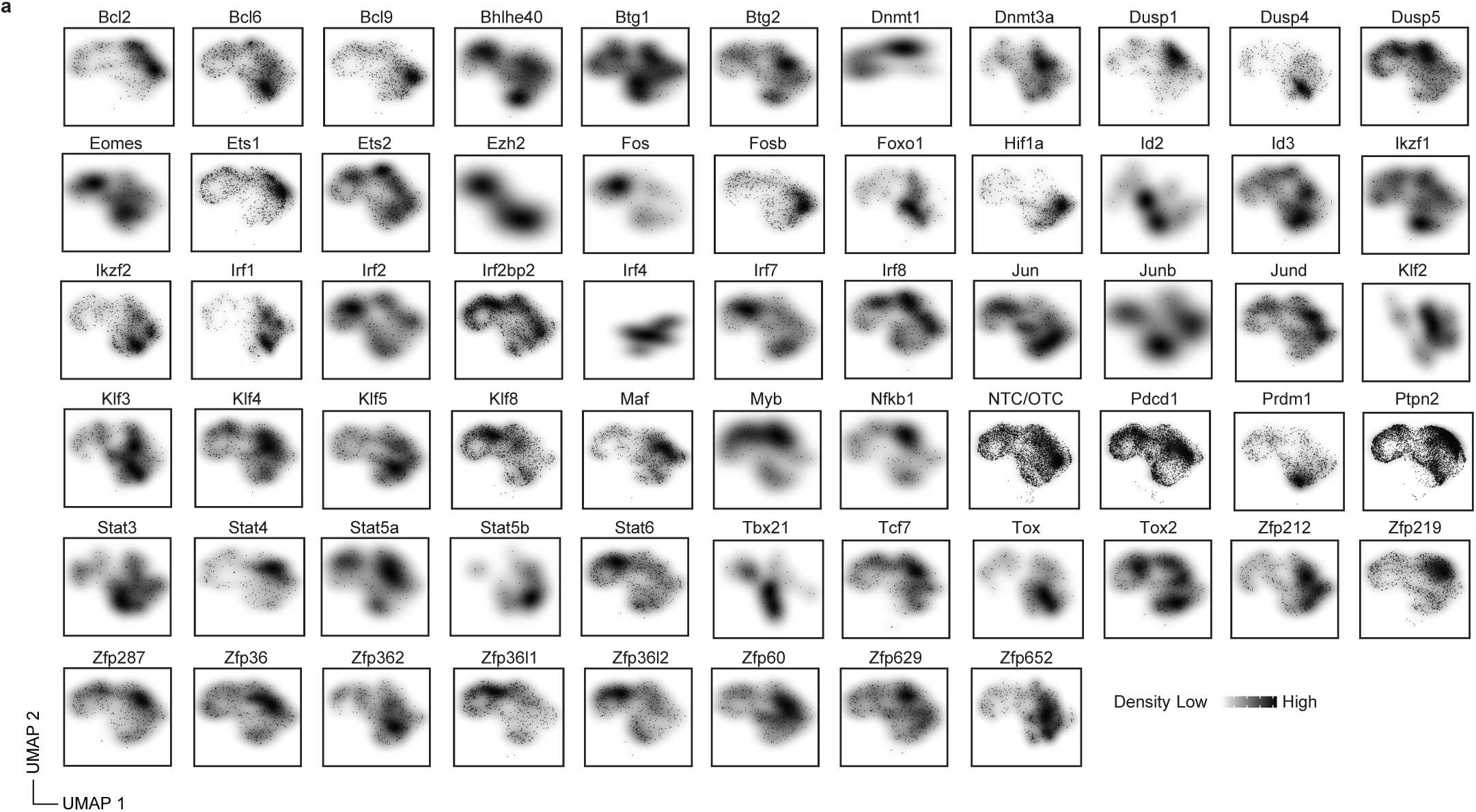
Cas9^+^ OT-1 CD8^+^ T cell density shifts by each gene perturbation in young vs. aged tumors, related to Figure 2. **a**, Cell density projections for various T cell clusters/subsets in tumors based on gene targets. Lighter colors indicate low cell density, while darker colors show high cell density.

**Extended Data Fig. 5.**
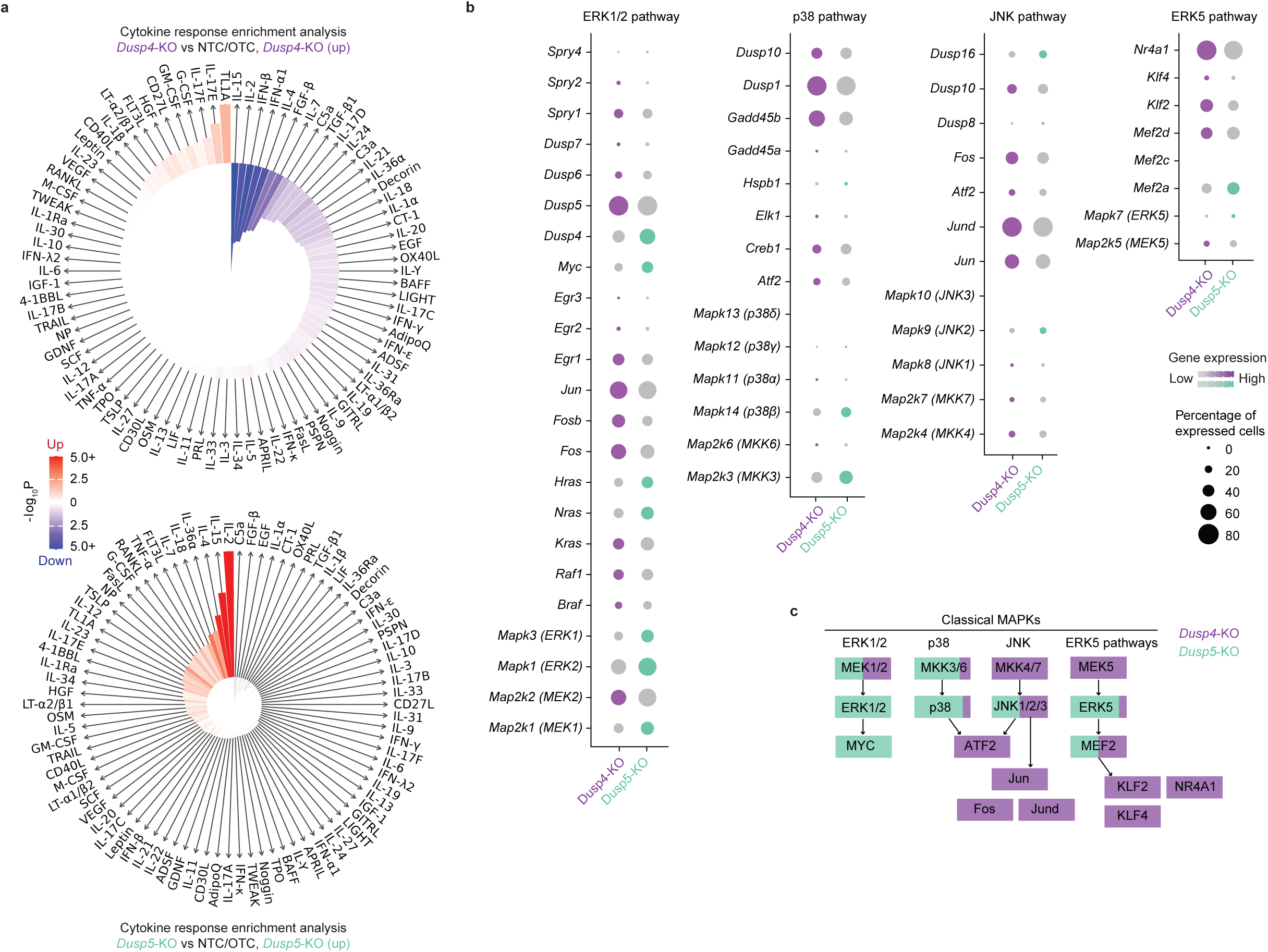
Distinct transcriptional profiles between *Dusp4*-KO and *Dusp5*-KO T cells, related to Figure 2. **a**, Immune Response Enrichment Analysis (IREA) cytokine enrichment plot showing the enrichment score (ES) for each of the 86 cytokine responses in CD8^+^ cells after *Dusp4* or *Dusp5* knockout (KO). Bar length represents the ES, shading represents the FDR-adjusted P value (two-sided Wilcoxon rank-sum test), with darker colors representing more significant enrichment (red, enriched in KO; blue, enriched in NTC/OTC). The upper plot is *Dusp4*-KO, and the lower plot is *Dusp5*-KO. **b-c**, Dotplot displaying upregulated genes in *Dusp4*-KO (purple) and *Dusp5*-KO (light-green) related to the mitogen-activated protein kinase (MAPK) pathway. Circle size reflects the percentage of expressed cells, and color intensity indicates gene expression levels, with darker shades representing higher expression.

**Extended Data Fig. 6.**
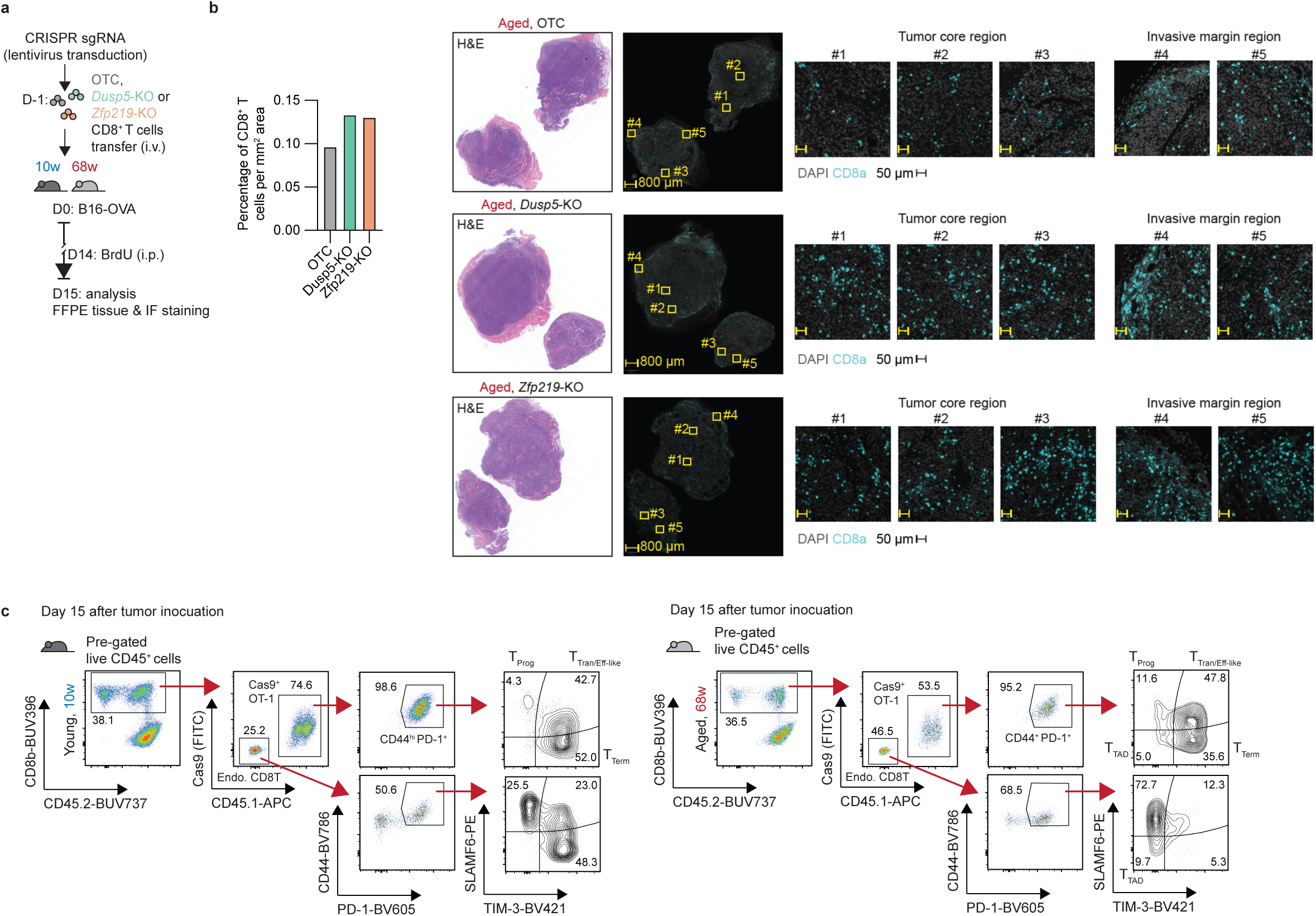
*Dusp5* and *Zfp219* KO enhances tumor control, related to Figure 3. **a-b**, Schema of the experimental design for aged B16-OVA tumor-bearing mice receiving *Dusp5*-KO, *Zfp219*-KO, or OTC control OT-1 T cells. Hematoxylin and eosin (H&E, middle) staining and immunofluorescence (IF, right) staining of tumor sections (left), with quantification of cell density (left). Representative CD8^+^ T cells (aqua) identified as CD8^+^ cells. **c**, Representative flow plots of CD8^+^ T cell subsets in tumors of young and aged mice. These include progenitor exhausted (T_Prog_, SLAMF6^+^ TIM-3^−^), transitory exhausted/effector-like (T_Tran_/_Eff-like_, SLAMF6^+^ TIM-3^+^), terminally exhausted (T_Term_, SLAMF6^−^TIM-3^+^), and tumor-infiltrating age-associated dysfunctional (T_TAD_, SLAMF6^−^TIM-3^−^) cells.

**Extended Data Fig. 7.**
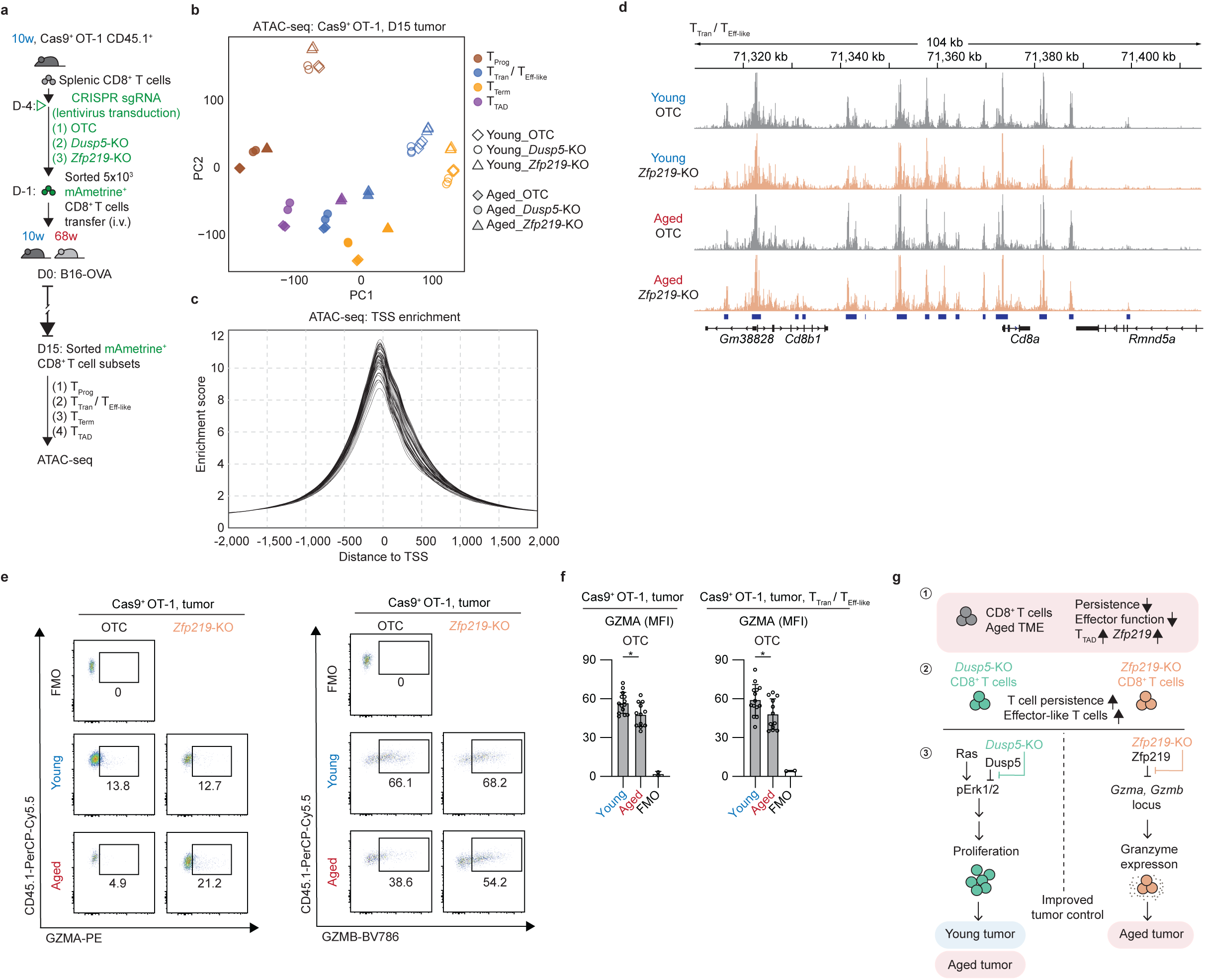
Epigenetic profiling of Cas9^+^ OT-1 CD8^+^ T cell subsets and granzyme expression after gene perturbation, related to Figure 4. **a,** Schema of the experimental design for isolating *Dusp5*-KO, *Zfp219*-KO, or OTC OT-1 T cells for ATAC-seq from young or aged B16-OVA tumor-bearing mice. **b-c**, Principal component analysis (PCA) projection and transcriptional start site (TSS) enrichment of chromatin accessibility profiles from tumor-infiltrating OT-1 T cell subsets. **d**, Representative ATAC-seq tracks showing OTC (gray) and *Zfp219*-KO (orange) OT-1 cells at specified gene loci. **e-f**, Representative flow cytometry plots showing GZMA and GZMB expression in OT-T cells from tumors, along with quantification of the geometric mean fluorescence intensity (MFI) of GZMA. **g**, Schema illustrating the different mechanisms of *Dusp5*-KO and *Zpf219*-KO. Mean ± s.d. For **f**, significance was calculated using a two-sided Student’s t-test. Asterisks used to indicate significance correspond to the following: N.S. (not significant, *P* > 0.05), \**P* ≤ 0.05, \*\**P* ≤ 0.01, and \*\*\**P* ≤ 0.001.

## Supplementary Tables

Supplementary Table 1. Antibody information

Supplementary Table 2. sgRNA library list of genes and sgRNA sequences

Supplementary Table 3. Human pan-cancer scRNA-seq patient and dataset information

Supplementary Table 4. T cell persistence by sgRNA abundance in Cas9 OT-1 T cells

Supplementary Table 5. T cell persistence by sgRNA abundance in Cas9 OT-1 T cells

Supplementary Table 6: Gene expression markers of each cell cluster of Cas9 OT-1 T cells in tumor

Supplementary Table 7. Differentially expressed genes between *Dusp4*-KO vs NTC/OTC

Supplementary Table 8. Differentially expressed genes between *Dusp5*-KO vs *Dusp4*-KO

Supplementary Table 9. Percentage of T cells subsets after gene perturbation within aged tumors

Supplementary Table 10: Gene expression markers of each cell cluster of Cas9 OT-1 T cells in tdLN

Supplementary Table 11. Differentially expressed genes between *Dusp5*-KO vs NTC/OTC

Supplementary Table 12. Differentially expressed genes between *Zfp219*-KO vs NTC/OTC

